# PI(3,5)P_2_ asymmetry during mitosis is essential for asymmetric vacuolar inheritance

**DOI:** 10.1101/2024.06.13.598808

**Authors:** Mariam Huda, Mukadder Koyuncu, Cansu Dilege, Ayse Koca Caydasi

**Affiliations:** Department of Molecular Biology and Genetics, Koç University, Istanbul, Turkey

**Keywords:** Asymmetric cell division, Aging, Atg18, Phosphatidylinositol 3,5-bisphosphate, Vacuole

## Abstract

Phosphatidylinositol 3,5-bisphosphate (PI(3,5)P_2_) is a low-abundance signaling lipid that plays crucial roles in various cellular processes, including endolysosomal system structure/function, stress response, and cell cycle regulation. PI(3,5)P_2_ synthesis increases in response to environmental stimuli, yet its behavior in cycling cells under basal conditions remained elusive. Here, we analyzed spatiotemporal changes in PI(3,5)P_2_ levels during the cell cycle of *S. cerevisiae.* We found that PI(3,5)P_2_ accumulates on the vacuole in the daughter-cell while it disappears from the vacuole in the mother-cell during mitosis. Concomitant with the changes in PI(3,5)P_2_ distribution, the daughter-vacuole became more acidic, whereas the acidity of the mother-vacuole decreased during mitosis. Our data further showed that both PI(3,5)P_2_ and the PI(3,5)P_2_ effector protein Atg18 are determinants of vacuolar-pH asymmetry and acidity. Our work, thus, identifies PI(3,5)P_2_ as a key factor for establishment of vacuolar-pH asymmetry, providing insights into how the mother cell ages while the daughter cell is rejuvenated.

## Introduction

Phosphorylated phosphatidylinositol lipids (PIPs) are signaling lipids that serve as master regulators of multiple cellular processes. Phosphatidylinositol-3,5-bisphosphate (PI(3,5)P_2_) constitutes one of the least abundant and least understood PIPs. Being a PIP of the endolysosomal system, PI(3,5)P_2_ plays critical roles in diverse aspects of cell biology such as membrane trafficking, stress response, vacuole/lysosome structure/function and cell cycle regulation (Han and Emr, 2011; Huda et al., 2023; Jin et al., 2014; Jin and Weisman, 2015; McCartney et al., 2014a; McCartney et al., 2014b; Rutherford et al., 2006).

PI(3,5)P_2_ is synthesized through phosphorylation of the phosphatidylinositol 3-phosphate (PI3P) by the PI3P-5kinase, PIKfyve in humans and Fab1 in yeast (Gary et al., 1998; Sbrissa et al., 1999). In human cells PIKfyve localizes at endolysosomal membranes (early, late, recycling endosomes and lysosomes), predominantly on recycling endosomes (Giridharan et al., 2022). In yeast, Fab1 predominantly decorates the membrane of the vacuole (yeast lysosomes) and localizes to some extend on signaling endosomes (Chen et al., 2021). On these membranes, PIKfyve/Fab1 resides within a protein complex composed of conserved proteins namely Vac14 (ArPIKfyve in humans) and Fig4 (Sac3 in humans) required for PIKfyve/Fab1 function (Barlow-Busch et al., 2023; Dove et al., 2002; Duex et al., 2006a; Lees et al., 2020). In yeast, additionally, a vacuolar membrane protein named Vac7 is part of the Fab1-complex and has an essential role in PI(3,5)P_2_ synthesis (Duex et al., 2006a; Duex et al., 2006b).

Under basal conditions, PI(3,5)P_2_ levels are scarce; however, PI(3,5)P_2_ abundance increases suddenly in response to environmental stimuli such as insulin treatment in mammalian cells and hyperosmotic shock in budding yeast (Bridges et al., 2012; Duex et al., 2006a; Jones et al., 1999; McCartney et al., 2014b; Sbrissa et al., 1999; Sbrissa and Shisheva, 2005; Tsujita et al., 2004). This is best observed in budding yeast, in which PI(3,5)P_2_ levels rise rapidly and transiently up to 20-fold upon hyperosmotic shock (Dove et al., 1997; Duex et al., 2006a). The abrupt response of PI(3,5)P_2_ synthesis to hyperosmotic shock made the budding yeast a great model to understand PI(3,5)P_2_ regulation and function under hypertonic conditions. A growing body of evidence, however, suggests that PI(3,5)P_2_ also functions in cycling cells under basal conditions (Huda et al., 2023; Jin et al., 2014; Jin et al., 2022; Jin and Weisman, 2015). Nevertheless, low levels of PI(3,5)P_2_ and lack of a well-established biosensor for in vivo detection of PI(3,5)P_2_ have been obstacles to the study of PI(3,5)P_2_ under isotonic conditions. In yeast, vacuolar localization of the PI(3,5)P_2_ binding protein Atg18, which is similar to WIPI proteins in mammals, was shown to correlate well with Fab1 activity (Dove et al., 2004; Efe et al., 2007; Gopaldass et al., 2017). Yet, Atg18 also binds to PI3P (Busse et al., 2015), and the Atg18 mutant that only binds PI(3,5)P_2_ but not to PI3P (Atg18-Sloop) has also reduced binding to PI(3,5)P_2_ (Gopaldass et al., 2017), challenging its use as an in vivo sensor. Recently, a new probe, SnxA from *Dictyostelium discoideum* was identified as an extremely selective PI(3,5)P_2_-binding protein (Vines et al., 2023). Its use as a PI(3,5)P_2_ reporter was characterized in mammalian cells and in *D. discoideum* (Vines et al., 2023), which now provided the means to study PI(3,5)P_2_ under basal conditions.

In this study, using SnxA and Atg18 as in vivo probes for PI(3,5)P_2_, we found that PI(3,5)P_2_ is asymmetrically present on yeast vacuoles during mitosis. PI(3,5)P_2_ accumulated on the membrane of the vacuole that resided in the daughter cell, while it disappeared from the vacuole that stayed in the mother cell. Asymmetry of PI(3,5)P_2_ was established shortly before anaphase onset, maintained during anaphase, and diminished with mitotic exit. PI(3,5)P_2_ precursor PI3P and Fab1-complex components were symmetrically present on the mother and daughter vacuoles, whereas expression of a hyperactive *FAB1* allele caused symmetric distribution of PI(3,5)P_2_ suggesting that asymmetric PI(3,5)P_2_ production is controlled at the level of Fab1 activity rather than the levels of the precursor or Fab1-complex proteins. Employing a ratiometric in vivo vacuolar-pH sensor, we observed a pH drop in the daughter vacuole, while it increased in the mother vacuole, concomitant with changes in PI(3,5)P_2_ distribution during mitosis. We further showed that the asymmetry of PI(3,5)P_2_ and the PI(3,5)P_2_ effector Atg18 determine the asymmetry of vacuolar pH.

## Results and Discussion

### Yeast codon optimized SnxA can be used as an in vivo sensor for PI(3,5)P_2_ in budding yeast

In order to monitor PI(3,5)P_2_ in budding yeast, we made use of the recently identified genetically coded PI(3,5)P_2_ sensor SnxA, which was tested in mammalian cells and in amoebae (Vines et al., 2023). To address whether SnxA would work as an in vivo PI(3,5)P_2_ sensor in budding yeast, first, we constructed yeast codon-optimized SnxA C-terminally fused with GFP and expressed under TEF1 promoter. Expression of SnxA-GFP integrated into the yeast genome did not alter the vacuole size or cell growth, indicating that SnxA expression does not interfere with PI(3,5)P_2_ function (Figure S1A-C). SnxA-GFP colocalized with the vacuolar membrane marker Vph1-3xmCherry in wildtype yeast cells, assessed by Pearson Correlation Coefficient (PCC) (Figure 1A) (Stauffer et al., 2018). Colocalization was lost in the absence of the PI 5-kinase *FAB1*, *VAC7* or *VAC14*, which are essential for PI(3,5)P_2_ synthesis (Figure 1A). Thus, we conclude that SnxA vacuole membrane localization is PI(3,5)P_2_ dependent in budding yeast. In line with this notion, SnxA co-localization with Vph1 increased in cells expressing a hyperactive Fab1 allele (*fab1-ha, fab1-14)* (Duex et al., 2006b), which leads to overproduction of PI(3,5)P_2_ (Figure 1B). Likewise, *fab-ha* expression fully rescued the loss of SnxA vacuole localization in *vac14*Δ cells and markedly increased SnxA colocalization with Vph1 in *vac7*Δ cells (Figure 1B), which is consistent with *fab-ha* rescue of PI(3,5)P_2_ levels in these cells (Duex et al., 2006b; Gary et al., 2002).

**Figure 1.**
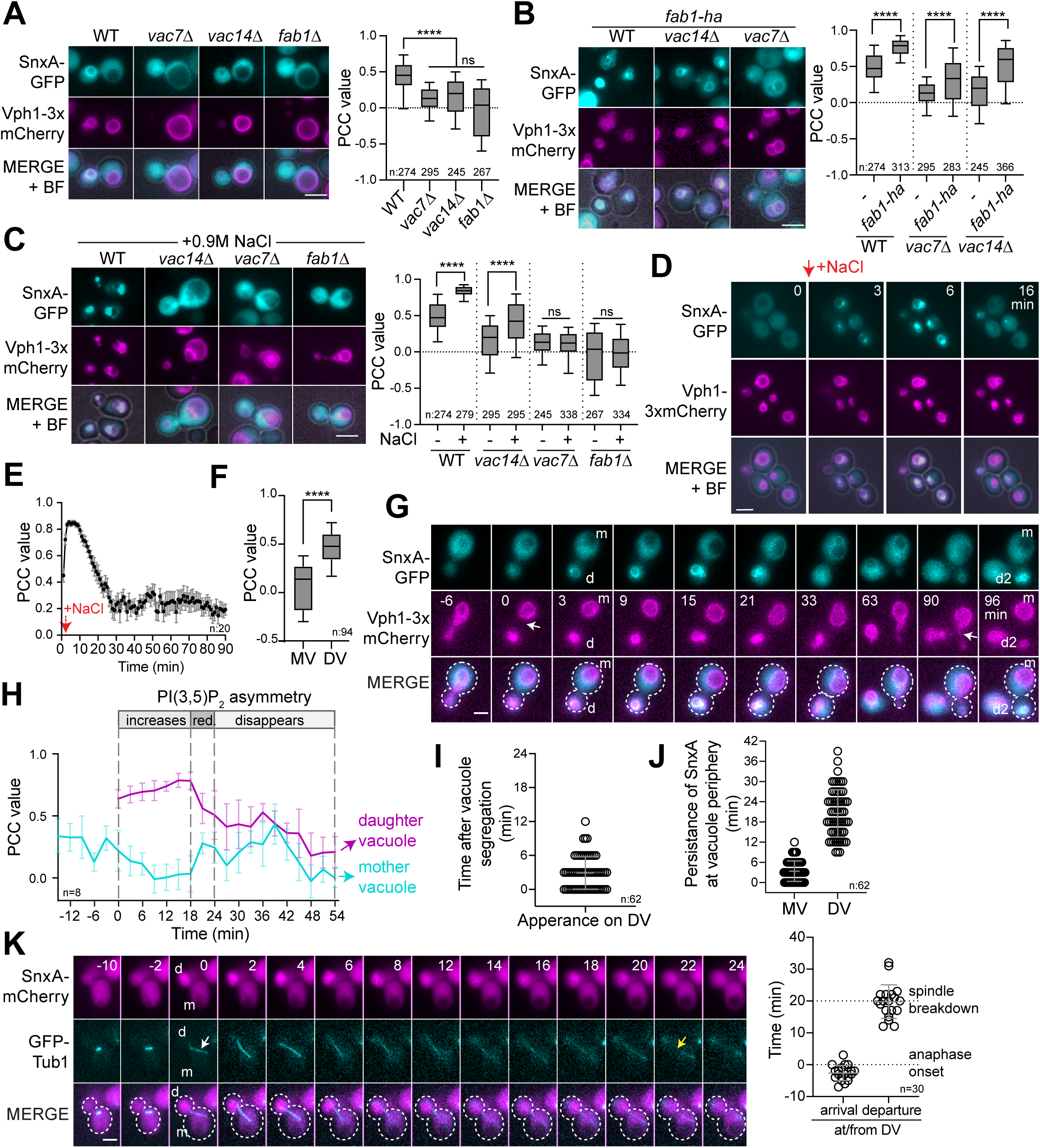
- PI(3,5)P_2_ sensor SnxA indicates daughter vacuole specific PI(3,5)P_2_ synthesis during late mitosis. A-C. Analysis of SnxA-GFP colocalization with the vacuolar marker Vph1-3xmCherry using still images of log-phase cell populations. Box-and-whisker plots of Pearson Correlation Coefficient (PCC) and representative still images are presented. PCC values were obtained using the EzColocalization plugin in ImageJ by selecting the vacuoles based on Vph1-3xmCherry as a vacuole marker (see materials methods for details). PCC provides a metric for likelihood of co-existence of SnxA-GFP and Vph1-3xmCherry signals in the given area, thus objectively assessing colocalization. Data shown are comparison of wildtype cells (WT) and PI(3,5)P_2_ deficient mutants (MHY286, MHY304, MHY302, and AAY007) in (A), comparison of these cells (-) with cells expressing *fab1-ha* (MHY286, MHY288, MHY304, MHY305, MHY302, and MHY303) in (B), and comparison of basal conditions (-) with hyperosmotic shock conditions (+) in (C). Experiments shown in A, B and C were performed altogether, thus the controls (WT, *vac7*Δ, *vac14*Δ and *fab1*Δ) are identical in all graphs. In box-and-whisker plots, boxes extend from the 25th to 75th percentiles. Lines inside the boxes are the median. Whiskers point to the 10^th^ and 90^th^ percentiles. Ordinary one-way ANOVA was performed with Fisher’s LSD test. ****: p<0.0001, ns: non-significant with p≥0.1. n: sample size. Data shown comes from four independent experiments. D-E. Analysis of the effect of hyperosmotic shock on SnxA-GFP vacuole periphery localization through time-lapse microscopy. Mean PCC values of SnxA-GFP and Vph1-3xmCherry during the time-lapse series are plotted (E) along with images from selected time points of representative cells (D). The graph represents the mean of PCC values obtained from 20 cells (MHY286) subjected to 0.9 M NaCl treatment. Error bars show the standard error of the mean. Red arrows indicate the time point of NaCl addition to the culture dish. Scale bars: 3 μm, BF: bright field. F. Box-and-whisker plots showing PCC values of SnxA-GFP colocalization with the vacuolar marker Vph1-3xmCherry on the mother vacuole (MV) and daughter vacuole (DV) in log-phase WT cells (MHY286) with segregated vacuoles. Boxes show the 25th to 75th percentiles, whiskers point to the 10^th^ and 90^th^ percentiles, and lines inside the boxes are the median. See Figure 1A for representative images. Unpaired two-tailed t-test was performed. ****: p<0.0001. Data shown comes from three independent experiments. G-H. Time-lapse microscopy analysis of the SnxA-GFP localization at the vacuole periphery during the cell cycle of WT cells (MHY286). Image series of selected time points from a representative cell is shown in (B). Time-dependent changes in PCC values of SnxA-GFP colocalization with the vacuolar marker Vph1-3xmCherry on the mother and daughter vacuoles are shown in the graph (C). The graph represents the mean of individual cells aligned with respect to the end of vacuole segregation (t=0). Error bars are the standard error of the mean. The white arrow in B indicates the completion of vacuole segregation (t=0). The periods in which PI(3,5)P_2_ asymmetry increases (increases), reduces (red.) and disappears are indicated on the graph. Data comes from two independent experiments. I-J. Analysis of SnxA-GFP localization on the mother and daughter vacuole periphery with respect to the timing of SnxA-GFP signal appearance on the daughter vacuole (DV) relative to the end of vacuole segregation (t=0) (I), and the duration of SnxA-GFP localization on the mother vacuole (MV) and daughter vacuole (DV) periphery (J). Circles on scatter plots indicate measurements of individual cells. Black lines in scatter blots are mean and standard deviation. Data comes from two independent experiments. K. Time-lapse microscopy-based analysis of SnxA-mCherry localization with respect to spindle elongation in WT cells containing GFP-Tub1 (MHY295). Selected time points from a representative cell are shown. White arrow marks the time point of anaphase onset (t=0) designated by the start of fast spindle elongation. Yellow arrow marks the timing of spindle breakdown as an indicative of mitotic exit. The graph shows the timing of SnxA-mCherry accumulation on the daughter vacuole (arrival at DV) with respect to the anaphase onset (t=0), and SnxA-mCherry disappearance from the daughter vacuole (departure from DV) with respect to the spindle breakdown (t=20). Scale bars: 3 μm. n: sample size. m: mother, d: daughter

It is well established by biochemical methods that the amount of PI(3,5)P_2_ increases drastically but transiently in response to hyperosmotic shock in budding yeast (Dove et al., 1997; Duex et al., 2006a) – such that within 5 minutes after hyperosmotic shock (0.9M NaCl treatment) PI(3,5)P_2_ levels increase ∼20 fold, elevated PI(3,5)P_2_ persists for ∼10 min, after which it starts decreasing, and restores near basal levels ∼30 minutes after initial exposure to hyperosmotic shock (Duex et al., 2006a). Hence, to capture the condition with high PI(3,5)P_2_ levels, we measured SnxA-GFP vacuole periphery localization ∼10 min after 0.9M NaCl treatment. SnxA colocalization with Vph1 increased dramatically upon hyperosmotic shock of logarithmically growing wildtype cells (Figure 1C). Salt treatment also caused an increase in SnxA-GFP vacuole periphery localization in *vac14*Δ cells but not in *vac7*Δ and *fab1*Δ (Figure1C), which aligns with previous data that hyperosmotic shock triggers PI(3,5)P_2_ production in *vac14*Δ cells, albeit at a low level, but not in *vac7*Δ and *fab1*Δ cells (Duex et al., 2006a). We further quantified SnxA vacuole localization upon hyperosmotic shock through time-lapse microscopy. Colocalization of SnxA-GFP with Vph1-3xmCherry prominently increased within the first 2 min following hyperosmotic shock (Figure 1D-E, Supplementary movie 1). High levels of SnxA colocalization with Vph1 were maintained for ∼7 min, after which it started decreasing (Figure 1D-E). Colocalization was diminished ∼30 min after the hyperosmotic shock (Figure 1D-E), in a similar manner measured by biochemical methods (Duex et al., 2006a).

We further observed that SnxA vacuole localization slightly differed in cells under basal conditions and under hyperosmotic shock as well as in cells expressing *fab1-ha*. While the wildtype cells under basal conditions had SnxA mostly homogenously decorating the vacuole periphery, upon hyperosmotic shock or *fab1-ha* expression SnxA-GFP became concentrated on vacuole-vacuole contact sites and/or fission sites (Contact/Fission, Figure S1D-E) or appeared more concentrated on some parts of the vacuole (patchy appearance, Figure S1D-E), supporting the role of PI(3,5)P_2_ in vacuole fission. Notably, under hyperosmotic shock conditions and in cells expressing *fab1-ha* we also noticed that ∼20% of the cells had strong SnxA-GFP signal in the vacuolar lumen in addition to the SnxA signal at vacuole periphery (Figure S1E-F), which may indicate additional presence of PI(3,5)P_2_ on vacuolar membranes that lacked the Vph1 signal or on other membranes (ie. autophagic vesicles).

Taken together, these data establish that SnxA can be used as an in vivo sensor for PI(3,5)P_2_ in budding yeast, similar to its usage in amoeba and mammalian cells (Vines et al., 2023).

### More PI(3,5)P_2_ is present on the daughter vacuole during yeast mitosis

While assessing SnxA-GFP colocalization with Vph1-3xmCherry, we noticed that in cells with segregated vacuoles, SnxA-GFP localization on the vacuole periphery was asymmetric, with a strong preference to localize on the vacuole that resides in the daughter cell compartment (daughter vacuole) (Figure 1A and 1F). On the vacuole of the mother cells (mother vacuole), however, SnxA-GFP was barely detectable (Figure 1A and 1F).

Next, we asked when SnxA asymmetry was established. To this end, we performed time-lapse microscopy of Vph1-3xmCherry SnxA-GFP containing cells. SnxA-GFP signal was low or undetectable on the vacuole during vacuole segregation to the daughter cell (Figure 1G-H). Within 3 minutes after completion of vacuole segregation (t=0), SnxA-GFP accumulated on the daughter vacuole (Figure 1I, Supplementary movie 2). SnxA-GFP signal persisted for ∼21 min at the daughter vacuole, after which the signal was greatly reduced (Figure 1J). Quantification of SnxA vacuole periphery localization dynamics revealed that SnxA asymmetry increased for ∼15-18 min after vacuole segregation (Figures 1G-H). During this time SnxA localization on the daughter vacuole increased, whereas it was completely diminished from the mother vacuole (Figures 1G-H). After this period, SnxA localization to the daughter vacuole started to decrease, concomitant with a slight increase of SnxA localization on the mother vacuole, causing a decrease in the SnxA asymmetry (Figures 1G-H, Supplementary movie 2). Finally, SnxA localized weakly but equally on both vacuoles, reaching a nearly symmetric state (Figures 1G-H). Using GFP-Tub1 as a mitotic marker, we next addressed the timing of SnxA accumulation on the daughter vacuole with respect to the anaphase onset and mitotic exit. SnxA-3mCherry appeared on the daughter vacuole 3±2 min (mean±SD) before anaphase onset and disappeared near mitotic exit (±5 min) (Figure 1K, Supplementary movie 3).

We thus conclude that PI(3,5)P_2_ is indeed synthesized in cycling cells under basal conditions. It peaks on the daughter vacuole during mitosis, whereas it disappears from the mother vacuole, creating an asymmetry. PI(3,5)P_2_ asymmetry is interrupted with mitotic exit.

### Atg18 localization at the vacuole periphery correlates with PI(3,5)P_2_ levels and mimics SnxA localization

As an alternative way to measure PI(3,5)P_2_ levels on vacuolar membranes in living cells, we next analyzed the localization of Atg18, a known protein that binds PI(3,5)P_2_ with higher affinity, and PI3P with a lower affinity (Dove et al., 2004; Efe et al., 2007; Gopaldass et al., 2017; Takatori et al., 2016). PI(3,5)P_2_ binding pool of Atg18 associates with the vacuolar membrane (Gopaldass et al., 2017). In concordance with this, wildtype Atg18-mNeongreen (Atg18-mNG) colocalized with Vph1-3xmCherry in logarithmically growing cells upon hyperosmotic shock (Figure S1G). Atg18-SlooP mutant, which binds PI(3,5)P_2_ but not PI3P (Gopaldass et al., 2017), also colocalized with Vph1; whereas the Atg18-FGGG mutant that does not bind PI(3,5)P_2_ and PI3P (Gopaldass et al., 2017) failed to localize on the vacuolar membrane (Figure S1G). Atg18-mNG also colocalized with Vph1 under basal conditions (Figure S1H). Hyperosmotic shock or expression of *fab1-ha* increased colocalization of Atg18-mNG with Vph1 (Figure S1H) whereas, deletion of *VAC7* or *VAC14* diminished Atg18 vacuole periphery localization (Figure S1H). Similar to the results obtained using SnxA, *fab1-ha* rescued vacuole localization of Atg18 in both *vac7*Δ and *vac14*Δ cells, whereas salt treatment only rescued vacuole localization in *vac14*Δ (Figure S1H). Accordingly, we conclude that Atg18 localization at the vacuole periphery is indicative of PI(3,5)P_2_ levels. Furthermore, time-lapse analysis of Atg18-mNG colocalization with Vph1-3xmCherry yielded similar results to that of SnxA (Figure 2). Atg18 explicitly localized to the vacuole that migrated to the daughter cell, but not to the vacuole that stayed in the mother cell in mitosis (Figure 2A-B, Supplementary movie 4) with similar timing as SnxA (Figure 2C-E). Thus, using two independent PI(3,5)P_2_ in vivo markers, Atg18 and SnxA, these results establish that PI(3,5)P_2_ is specifically present on the daughter vacuole of the yeast during mitosis, whereas it diminishes from the mother vacuole.

**Figure 2.**
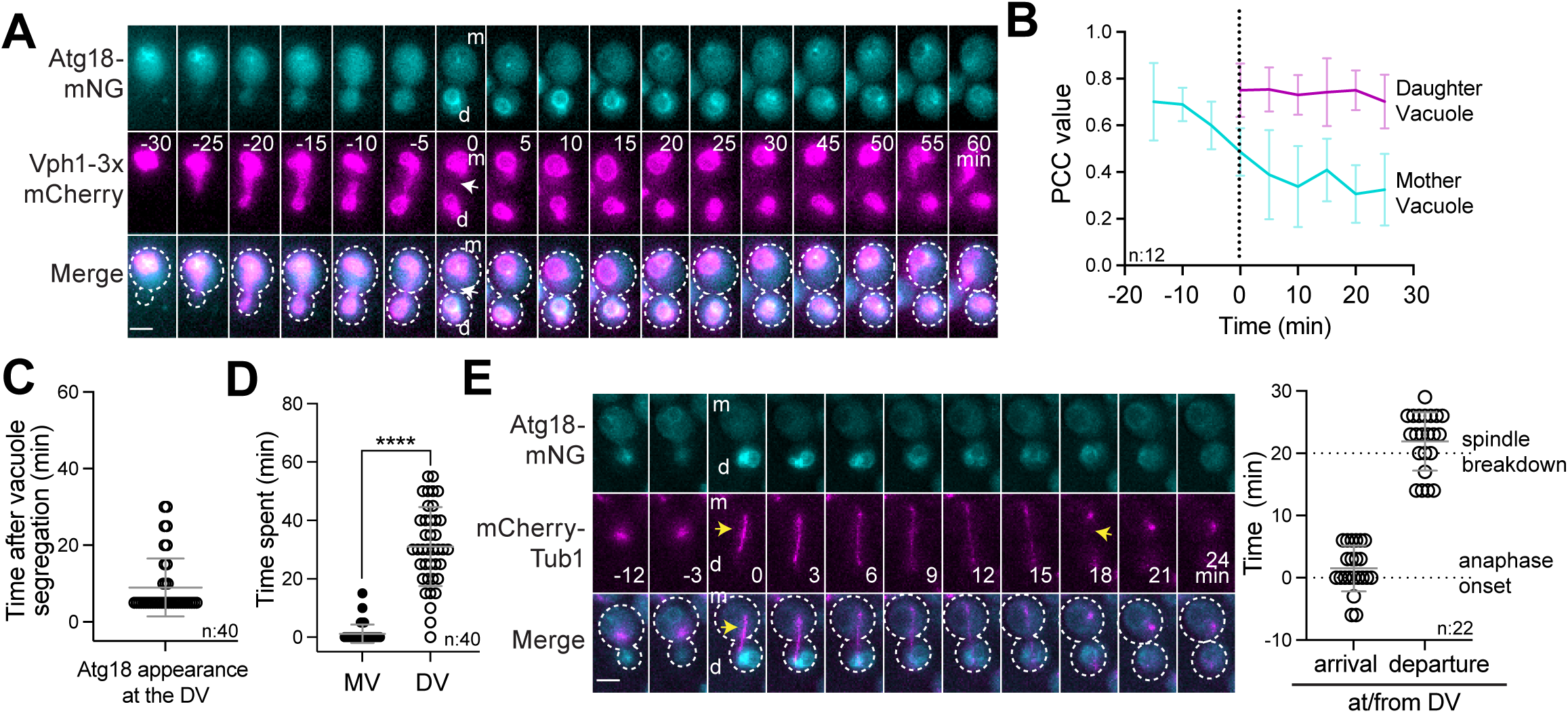
- Atg18-mNeongreen localizes predominantly to the daughter vacuole periphery. A-D. Time-lapse microscopy-based analysis of Atg18-mNeongreen (Atg18-mNG) vacuole periphery localization during the cell cycle in cells containing Vph1-3xmCherry (MUK089). Selected time points from a representative time-lapse series are shown (A) and mean PCC values of Atg18-mNG colocalization with Vph1-3xmCherry are plotted (B). Data is aligned with respect to the timing of vacuole segregation (t=0) in B and the error bars show the standard deviation. White arrow in A indicates the end of vacuole segregation (t=0). Timing of Atg18-mNG appearance on the daughter vacuole with respect to completion of vacuole segregation (C), and persistence of Atg18-mNG signal on both vacuoles (D) are plotted. Ordinary one-way ANOVA was performed with Fisher’s LSD test. **** p<0.0001. Data comes from 3 independent timelapse experiments. E. Atg18-mNG daughter vacuole localization with respect to the timing of anaphase onset and spindle breakdown in mCherry-Tub1 containing cells (MUK116). Time-lapse series are of a representative cell are shown. Yellow arrows mark the start of anaphase onset and spindle breakdown respectively. The graph shows Atg18-mNG signal appearance (arrival) and disappearance from the daughter vacuole (departure) relative to the timing of anaphase onset and spindle breakdown, respectively. Data comes from 2 independent timelapse experiments. Scale bars are 3 μm, n: sample size, DV: daughter vacuole, MV: mother vacuole. Circles in scatter plots are values of individual cells. Black lines in scatter plots represent the mean and standard deviation.

### Levels of PI3P and Fab1-complex components are symmetrically distributed on the mother and daughter vacuole

We reasoned that the asymmetry of PI(3,5)P_2_ may stem from differential levels of its precursor, PI3P, on mother and daughter vacuoles. To understand whether this is the case, we monitored PI3P in living cells using FYVE domain of mammalian Hrs (Gillooly et al., 2000; Obara et al., 2008). Unlike PI(3,5)P_2_, GFP-2xFYVE localized equally on mother and daughter vacuoles (Figure 3A). Thus, PI(3,5)P_2_ asymmetry is unlikely to be due to PI3P levels. Alternatively, PI(3,5)P_2_ asymmetry may stem from asymmetric distribution of Fab1, Vac7, and Vac14 proteins, which altogether constitute the Fab1-complex. Nevertheless, unlike Atg18, neither of these proteins were asymmetrically distributed on mother and daughter vacuoles (Figure 3B), suggesting that it is the activity of the Fab1 rather than the levels of Fab1-complex proteins on vacuoles that determine the asymmetry of PI(3,5)P_2_. Supporting this hypothesis, expression of the hyperactive form of Fab1 (*fab1-ha*) resulted in not only an increment of PI(3,5)P_2_ levels but also promoted equal levels of PI(3,5)P_2_ on both vacuoles, breaking the PI(3,5)P_2_ asymmetry (Figure 3C). Similarly, elevation of Fab1 activity by deletion of *ATG18* (Efe et al., 2007), also disrupted PI(3,5)P_2_ asymmetry (Figure 3C). These results altogether support that mechanisms establishing PI(3,5)P_2_ asymmetry is likely to regulate Fab1-complex activity, but not the abundance of Fab1-complex proteins on the vacuole periphery or the precursor PI3P.

**Figure 3.**
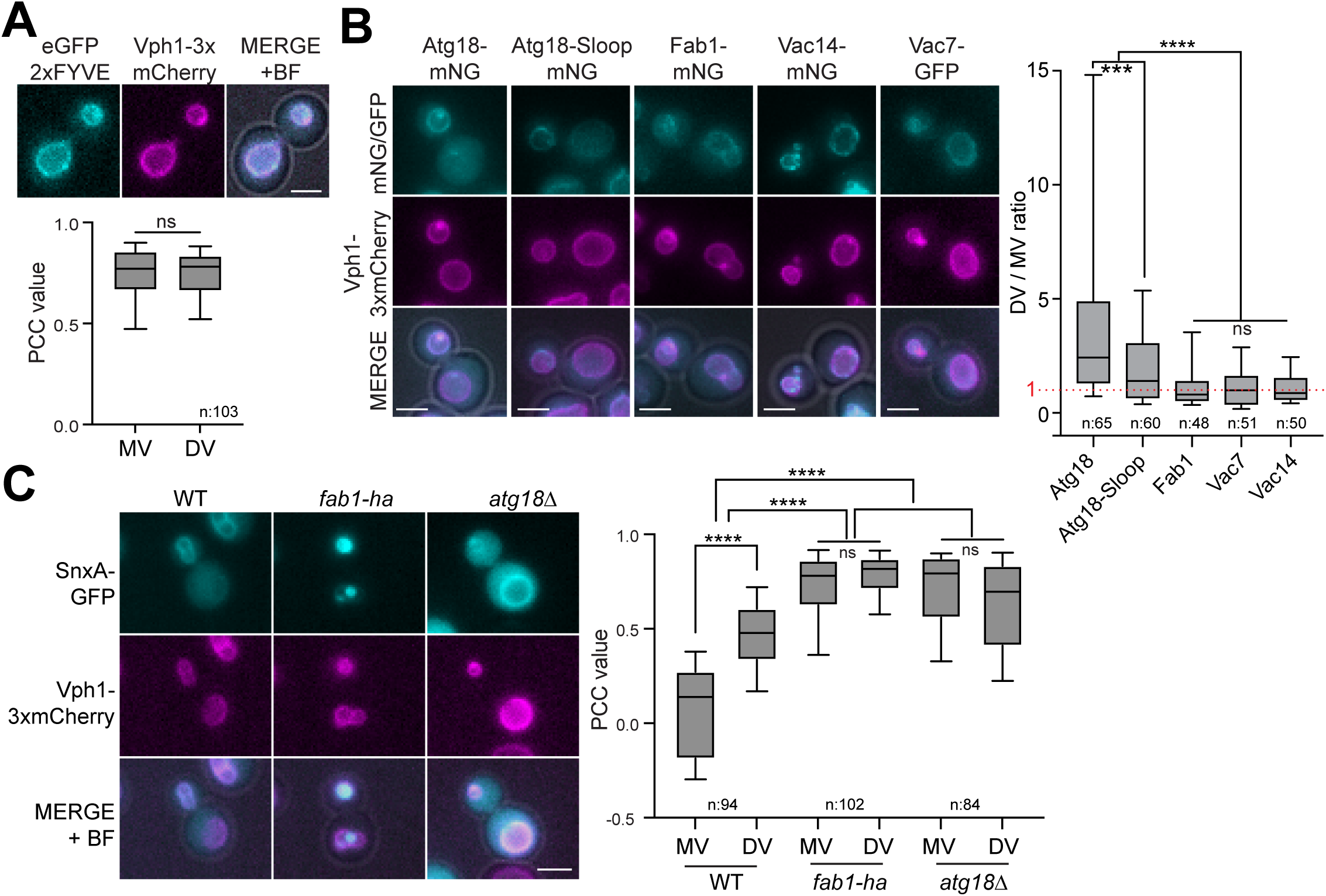
- PI3P and Fab1-complex components localize symmetrically on the vacuole periphery. A. Representative still images and PCC values showing EGFP-2xFYVE colocalization with the vacuolar marker Vph1-3xmCherry in WT cells (MHY252). Unpaired two-tailed t-test was performed. n.s: non-significant with p=0.9036. Data shown is from 2 independent experiments. B. Representative still images and daughter vacuole to mother vacuole (DV/MV) ratio of Atg18, Atg18-Sloop, Fab1, Vac14 and Vac7 (MUK089, MUK091, SGY008, SGY009, and MUK117) are shown. The red dashed line marks the DV/MV value of 1, which indicates symmetric distribution on both vacuoles without a preference for a specific vacuole. See materials methods for details. C. Representative still images and PCC values showing SnxA-GFP colocalization with Vph1-3xmCherry on the mother and daughter vacuoles of wildtype (WT) cells and *fab1-ha* or *atg18*Δ mutants (MHY286, MHY288, and MHY289). Data shown is from 2 independent experiments. In the box-and-whisker graphs, boxes show the 25th to 75th percentiles, whiskers show the 10^th^ and 90^th^ percentiles; lines inside the boxes are the mean. In B and C, ordinary one-way ANOVA was performed with Fisher’s LSD test. ***: p=0.0002, **** p<0.0001, ns: non-significant with p≥0.7441. Scale bars are 3 μm. BF: bright field.

### Asymmetry of vacuolar pH is established concomitant with PI(3,5)P_2_ asymmetry

Facilitating the assembly and activity of Vph1 containing vacuolar H^+^-ATPase (V-ATPase), PI(3,5)P_2_ functions in vacuole acidification (Banerjee et al., 2019; Li et al., 2014). Of importance, vacuolar pH also exhibits an asymmetric behavior between mother and daughter cells, such that the daughter vacuole is more acidic than the mother vacuole already before cytokinesis (Henderson et al., 2014; Hughes and Gottschling, 2012). We thus asked whether daughter vacuole acidification coincides with PI(3,5)P_2_ accumulation on the daughter vacuole. To assess vacuolar pH we employed the vSEP-mCherry ratiometric pH sensor in which, the vacuole-targeted acid-sensitive fluorophore vSEP is fused with pH-insensitive fluorophore mCherry (Okreglak et al., 2023). Consistent with the previous studies, vacuolar fluorescence intensity of vSEP increased linearly from pH 5 to 8 (Okreglak et al., 2023) (Figure S2A), whereas the mean fluorescence intensity of mCherry remained nearly constant within this pH range (Figure S2B) (Botman et al., 2019). Accordingly, vSEP/mCherry ratios exhibited a linear correlation in the 5-to-8 pH interval (Figure S2C), allowing usage of the vSEP/mCherry ratios in our experiments to evaluate vacuolar pH in this range.

To address the timing of daughter vacuole acidification, we performed time-lapse microscopy of cells carrying vSEP/mCherry. We quantified vSEP/mCherry ratios in daughter and mother vacuoles separately, and with respect to the timing of vacuole segregation (Figure 4A, Supplementary movie 5). Using our previous data that established dynamic changes of PI(3,5)P_2_ as a reference (Figure 1H), we compared the changes in vSEP/mCherry ratio with the changes in PI(3,5)P_2_ asymmetry. Our results showed that, shortly after the establishment of PI(3,5)P_2_ asymmetry, the pH of the daughter vacuole started to decrease rapidly, while the pH of the mother vacuole started to increase (Figure 4A). This suggests that acidification of the daughter vacuole follows elevation of PI(3,5)P_2_ levels on the daughter vacuole, whereas reduction of acidity of the mother vacuole follows the diminishment of PI(3,5)P_2_ on the mother vacuole. Supporting this view, concomitant with the reappearance of PI(3,5)P_2_ on the mother vacuole (Figure 1H, 18-24 min), the mother vacuole started to regain its acidity (Figure 4A, 18-24 min). Upon establishment of symmetric PI(3,5)P_2_, both mother and daughter vacuole pH exhibited similar behavior over time. However, the daughter vacuole remained more acidic than the mother vacuole. Our data thus shows that the vacuolar pH is established soon after vacuole segregation and further indicates that changes in PI(3,5)P_2_ asymmetry correlate with changes in the vacuolar pH.

**Figure 4.**
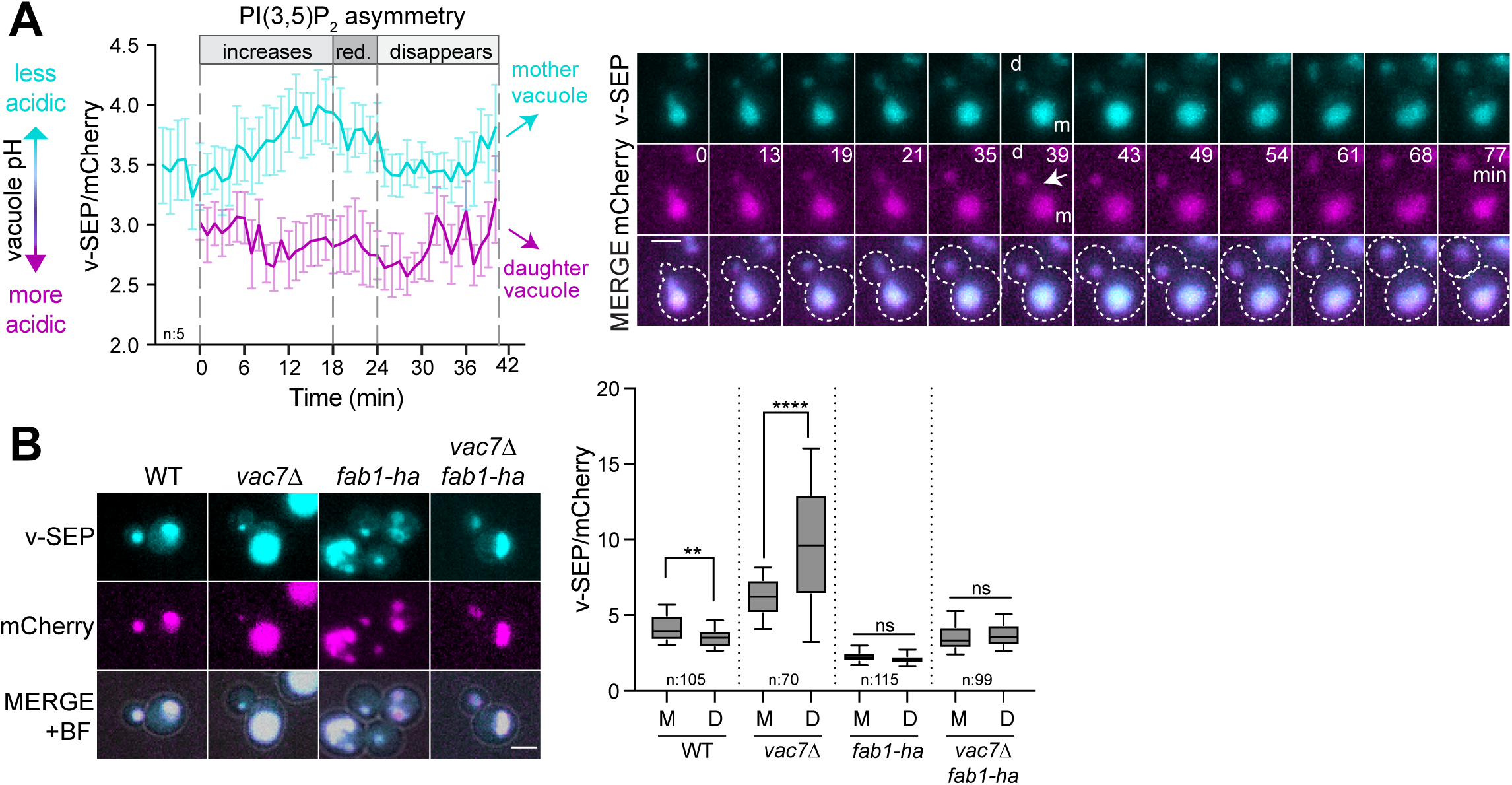
- Asymmetry of PI(3,5)P_2_ is critical for asymmetry of vacuolar pH. A. Assessing relative acidity of mother and daughter vacuoles during the cell cycle of wildtype cells (MHY283). v-SEP-to-mCherry ratios (v-SEP/mCherry) of daughter and mother vacuoles serve as an in vivo pH sensor. The graph shows mean v-SEP/mCherry ratios of five cells. Error bars are standard error of the mean. Cells are aligned with respect to the end of vacuole segregation (t=0). Selected time points from the time series of a representative cell are shown. White arrow marks the time of vacuole segregation (t=0). The periods in which PI(3,5)P_2_ asymmetry increases (increases), reduces (red.), and disappears are indicated on the graph based on the Figure 2. Data comes from 2 independent experiments. Scale bars: 3 μm. B. Representative still images and box-and-whisker plots of v-SEP/mCherry ratios between mother and daughter vacuole in WT (MHY283), *vac7*Δ (MHY291), *fab1-ha* (MHY257), and *vac7Δ fab1-ha* (MHY296). Data shown is from 2 independent experiments. Boxes show 25th and 75th percentiles, whiskers indicate 10^th^ and 90^th^ percentiles, the line in the boxes represents the median. Ordinary one-way ANOVA was performed with Fisher’s LSD test. **** p<0.0001, ** p=0.0087, ns: non-significant with p≥0.6617. Scale bar:3 μm, BF: bright field.

### PI(3,5)P_2_ and its asymmetry are crucial for the asymmetry of vacuolar pH

In order to understand whether PI(3,5)P_2_ and its asymmetry are required for vacuole acidity and asymmetry of the vacuolar pH, we quantified vSEP/mCherry ratios of daughter and mother vacuoles in the absence of PI(3,5)P_2_ (using *vac7*Δ cells) and under conditions that yield symmetric PI(3,5)P_2_ distribution (using *fab1-ha* expressing cells). Mother vacuole of wildtype cells had higher vSEP/mCherry ratios than the daughter vacuoles (Figure 4B), consistent with previous reports (Henderson et al., 2014; Hughes and Gottschling, 2012) and our time-lapse data (Figure 4A). Deletion of *VAC7* caused a remarkable increase in the pH of both vacuoles supporting the function of PI(3,5)P_2_ in vacuole acidity (Figure 4B). Intriguingly, *VAC7* deletion also resulted in relatively less acidic daughter vacuoles than the mother vacuoles, a situation opposing wildtype cells. *fab1-ha* expression, however, caused more acidic vacuoles and symmetric distribution of pH among the daughter and mother vacuoles in wildtype cells (Figure 4B). Furthermore, *vac7*Δ cells expressing *fab1-ha* had symmetric vacuolar pH, while the overall vacuolar acidity resembled that of the daughter vacuole of the wildtype cells (Figure 5B). These data altogether support that PI(3,5)P_2_ is required for vacuole acidification and further suggest that asymmetry of PI(3,5)P_2_ is crucial for asymmetry of vacuolar pH.

**Figure 5.**
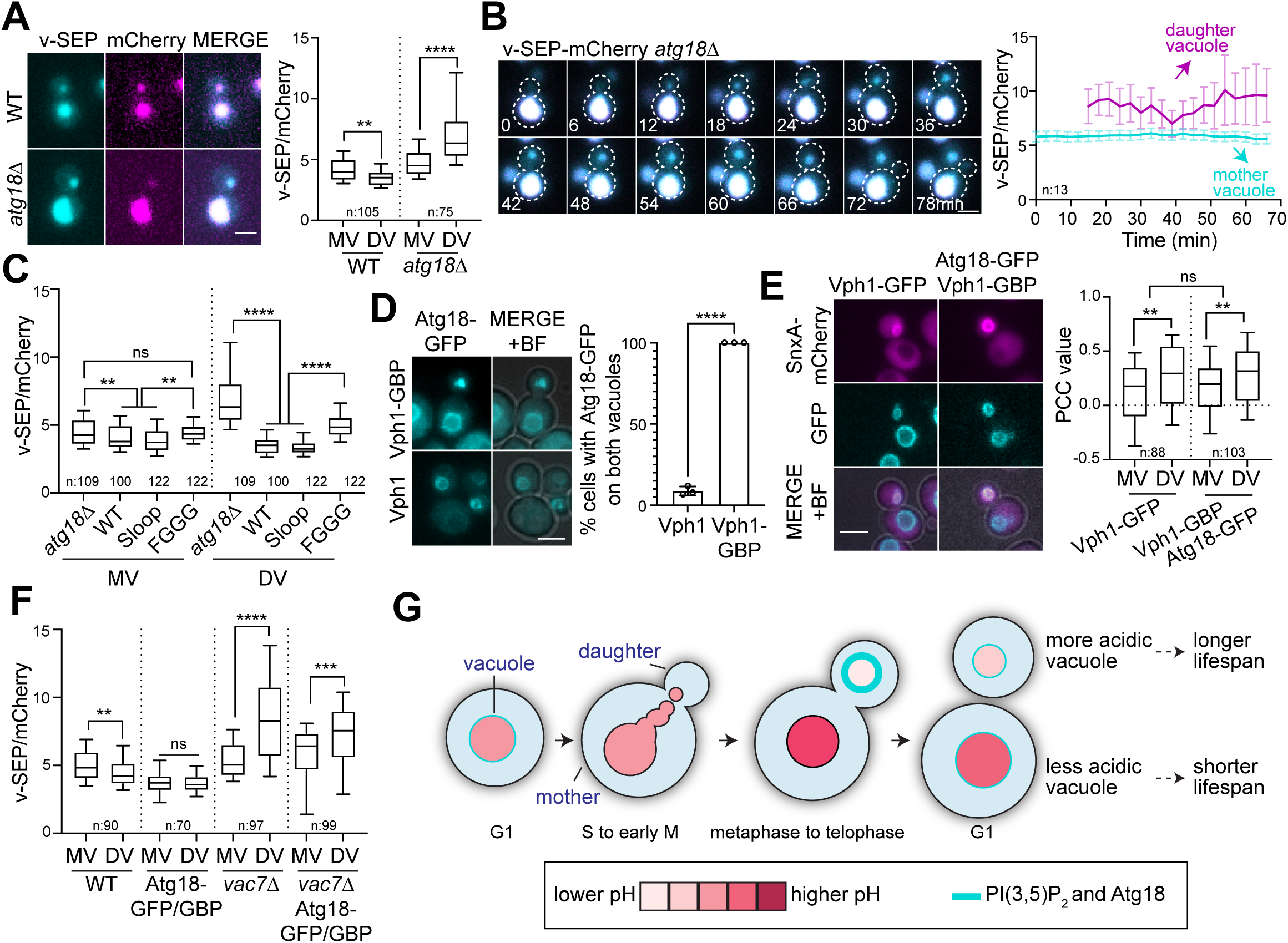
- Vacuole acidity depends on Atg18 and its binding to PI3,5P2. A. Representative still images and v-SEP/mCherry ratios of mother and daughter vacuoles in WT (MHY283) and *atg18*Δ (MHY277) cells. Data shown is from 2 independent experiments. B. Analysis of vacuolar pH via time-lapse microscopy in *atg18*Δ cells (MHY277). Selected time points from the time series are shown. The graph represents the mean of v-SEP/mCherry ratios of mother and daughter vacuoles as a function of time. Data from individual cells are aligned with respect to the vacuole segregation. Error bars show the standard error of the mean. C. Analysis of vacuolar pH in log-phase culture of WT (MHY283), *atg18*Δ (MHY277), Atg18-Sloop (MHY315), and Atg18-FGGG (MHY317) cells. Data comes from 2 independent experiments. D. Atg18-GFP vacuolar localization in cells with and without Vph1-GBP. Representative still images and the percentage of cells in which Atg18-GFP localizes on both vacuoles are shown (MHY223, and SEY082). Graph shows mean of three independent experiments. ∼100 cells were counted per strain per experiment. Error bars: standard deviation. ****: p<0.0001 according to unpaired two-tailed t-test. E. Analysis of SnxA-mCherry vacuole periphery localization in cells carrying Vph1-GFP and Atg18-GFP Vph1-GBP (MHY285, and MHY297). F. Effect of constitutive Atg18 targeting to both vacuoles (Atg18-GFP/GBP) on vacuolar pH (MHY283, MHY297, MHY291, and MHY312). Data shown comes from 2 independent experiments. In box-and-whisker graphs, boxes show the 25th and 75th percentiles, whiskers indicate the 10^th^ and 90^th^ percentiles, and the line in the boxes represents the median. In A, C, E and F, ordinary one-way ANOVA was performed with Fisher’s LSD test. **** p<0.0001, ** p≤0.0093, *** p=0.0004, and ns: non-significant with p≥0.6580. Scale bars: 3 μm. BF: bright field. G. Working model. PI(3,5)P_2_ accumulates on the vacuoles of the daughter cell while it is depleted from the vacuoles of the mother cell during mid-to-late mitosis. Concomitantly, daughter cell’s vacuole becomes acidified, and acidity of the mother cell’s vacuole declines. Thus, presence of PI(3,5)P_2_ on the daughter vacuole plays a role in daughter cell’s rejuvenation and absence of PI(3,5)P_2_ from the mother vacuole leads to mother cell’s aging.

### PI(3,5)P_2_-mediated vacuole acidification largely depends on Atg18

Next, we analyzed the vacuolar pH in *atg18*Δ cells in which PI(3,5)P_2_ appears symmetric on the daughter and mother vacuoles (Figure 3C). However, unexpectedly, we observed that *atg18*Δ cells had overall more alkalinized vacuoles, and similar to *vac7*Δ cells, the asymmetry of vacuolar pH was reversed such that the mother vacuoles were more acidic than the daughter vacuoles (Figure 5A). Analysis of vacuolar pH in *atg18*Δ cells through time-lapse microscopy confirmed the reversed asymmetry of the vacuolar pH (Figure 5B, Supplementary movie 6). Furthermore, vacuolar pH dynamics in the mother appeared more stable during mitosis in *atg18*Δ cells compared to the wildtype cells (Figure 5B), which is in support of a previous report (Okreglak et al., 2023). Thus, Atg18 is critical for vacuole acidity as well as asymmetry and dynamic aspects of vacuolar pH.

Vacuole acidity of *atg18*Δ cells was fully rescued by the Atg18-Sloop mutant that localizes to the vacuole membrane through binding to PI(3,5)P_2_ (Figure 5C). Introduction of the Atg18-FGGG mutant that cannot bind membranes, however, only partially rescued vacuolar pH (Figure 5C), suggesting that Atg18 vacuole periphery localization is necessary for vacuole acidity. Considering that Atg18 vacuole periphery localization requires its binding to PI(3,5)P_2_, we propose that Atg18 may have a role downstream of PI(3,5)P_2_ in vacuole acidification. We reasoned that if this were the case, artificial recruitment of Atg18 on the membrane of both vacuoles would lead to mother vacuole acidification and symmetric pH. In order to target Atg18 on both mother and daughter vacuoles, we tagged endogenous Vph1 and Atg18 at their C-terminus with GFP-binding protein (GBP) and GFP, respectively (Caydasi et al., 2017). This strategy successfully provided symmetric localization of Atg18 on the periphery of both vacuoles (Figure 5D). Constitutive presence of Atg18 on both vacuoles did not alter PI(3,5)P_2_ asymmetry (Figure 5E), yet caused more acidic and symmetric vacuolar pH (Figure 5F, WT vs Atg18-GFP/GBP). This data supports the role of vacuole membrane localized Atg18 in vacuole acidification. However, PI(3,5)P_2_ presence was essential for this role, as constitutive targeting of Atg18 to the vacuolar membranes in *vac7*Δ cells had only a minor effect on vacuole acidity (Figure 5F, *vac7*Δ vs *vac7*Δ Atg18-GFP/GBP). We do not yet know how Atg18 promotes vacuolar acidity. While it may interact with PI(3,5)P_2_ to support V-ATPase activity, Atg18 has not been shown to bind V-ATPase subunits. Given its role in transporting vacuolar enzymes (Harding et al., 1996; Yorimitsu and Klionsky, 2005), Atg18’s involvement in vacuole acidification may be also related to vacuole biosynthesis and maturation. If this is the case, it likely does not involve Vph1 sorting (Finnigan et al., 2012), as Vph1 localization isn’t affected by *ATG18* deletion (Figure 3C). Atg18’s role in vacuole acidification may also be linked to autophagy, which is required for vacuole acidification (Ruckenstuhl et al., 2014). Further research is needed to clarify this connection.

### PI(3,5)P_2_ production on the daughter vacuole is dependent on carbon sources but not on Pho85 and amino acid availability

Carbon sources are established regulators of vacuolar pH (Kane, 2016). To address whether they also regulate PI(3,5)P_2_ levels, we analyzed SnxA signal upon carbon source withdrawal and in media containing respiratory carbon sources (Ethanol/Glycerol medium). In these media, strikingly, SnxA signal disappeared from both the mother and daughter vacuole peripheries (Figure S2D). This data is in line with the role of PI(3,5)P_2_ upstream of V-ATPase assembly (Banerjee et al., 2019; Li et al., 2014) and the disassembly of V-ATPase observed in these media (Kane, 1995; Parra and Kane, 1998). It further suggests that the availability of carbon sources and the respiration/fermentation state of the cell is critical for PI(3,5)P_2_ synthesis.

Next, we tested the contribution of amino acid availability to PI(3,5)P_2_ asymmetry in mitotic cells, as availability of some amino acids, such as tyrosine and phenylalanine but not histidine, were shown to influence pH oscillations of the mother vacuole (Okreglak et al., 2023). PI(3,5)P_2_ asymmetry was maintained independently of amino acid supplementation (Figure S2E). However, in comparison to the media supplemented with all amino acids, colocalization on the mother vacuole was slightly increased in minimal media supplemented with only histidine, which was rescued by addition of tyrosine or phenylalanine to the media (Figure S2E). This suggests that amino acid availability does not contribute to the increased PI(3,5)P_2_ levels on the daughter vacuole during mitosis, but may have a minor impact on the mother vacuole associated PI(3,5)P_2_ during mitosis. Analysis of the effect of Pho85 on SnxA asymmetry led to a similar conclusion (Figure S2F). The nutrient responsive cyclin dependent kinase Pho85 phosphorylates Fab1 and is essential for PI(3,5)P_2_ production upon hyperosmotic shock (Jin et al., 2014). Pho85 also greatly influences the cell cycle linked pH oscillations of the mother vacuole (Okreglak et al., 2023). When we analyzed SnxA-GFP localization in mitotic cells, we observed that SnxA localization on the mother vacuole was impaired in cells with segregated vacuoles (Figure S2F). This is in line with previous conclusions that the cell cycle dependent oscillation of the mother vacuole is dependent on Pho85 (Okreglak et al., 2023). SnxA daughter vacuole localization, however, did not depend on Pho85 (Figure S2F). Thus, our data suggests presence of a mechanism different than Pho85 dependent phosphorylation of Fab1 to activate the Fab1-complex on the daughter vacuole during mitosis.

Altogether, our work revealed that the vacuolar signaling lipid PI(3,5)P_2_ and its effector protein Atg18 are primarily present on the daughter vacuole during mid-to-late mitosis. Our data further showed that the asymmetry of PI(3,5)P_2_ and Atg18 is critical for the asymmetry of vacuolar pH, which is established in mitosis. Notably, a previous study that focused on the pH of mother vacuoles showed that the vacuole alkalinizes before cell division and re-acidifies as cells divide. This oscillation of vacuolar pH was also found to be dependent on PI(3,5)P_2_ and Atg18 among other proteins (Okreglak et al., 2023). Our results align with these findings, and further describe how the pH of the daughter vacuole changes during mitosis, as well as how the pH of both vacuoles behaves in relation to the levels of PI(3,5)P_2_-Atg18 on the vacuole. Thus, our work establishes a link between PI(3,5)P_2_-Atg18 asymmetry and vacuolar pH asymmetry. Given that the decline of vacuolar acidity is a key factor contributing to yeast aging (Henderson et al., 2014; Hughes and Gottschling, 2012), we propose that PI(3,5)P_2_ asymmetry, by ensuring that daughter cells have more acidic vacuoles while mother cells have less acidic vacuoles, is one of the mechanisms leading to the aging of the mother cell and the rejuvenation of the daughter cell through asymmetric regulation of vacuolar pH (Figure 5G).

## Methods

### Yeast methods, strains, and plasmids

All yeast strains used in this study are isogenic with S288C and are listed in Table S1. Basic yeast methods and growth media were as described previously (Sherman, 1991). All yeast strains were grown at 30°C, and in synthetic complete media that contained all amino acids and glucose (2%) (SC-media, Glu) (Caydasi and Pereira, 2017) unless otherwise stated. For carbon source withdrawal experiments, cells were brought to log-phase in SC-media and shifted to synthetic media without glucose and with all amino acids (NC) for 2.5 hours before imaging.

For experiments that analyzed cells in ethanol and glycerol containing media (EG), synthetic media that contained all amino acids but lacked glucose was supplemented with 2% glycerol and 3% ethanol, cells were brought to log-phase in this media. For amino acid depletion or supplementation experiments, MHY333 was brought to log-phase in SC-media, SC-media that lacked all amino acids but histidine (85 mg/l), or in amino acid add back media in which SC-media that lacked all amino acids was supplemented with 85 mg/l of tyrosine or phenylalanine in addition to histidine.

The cassette PCR-based gene editing method was used for gene deletion and C-terminal tagging of endogenous proteins (Janke et al., 2004; Knop et al., 1999). v-SEP-mCherry containing strains were obtained as described by (Okreglak et al., 2023). GFP-Tubulin, mCherry-Tubulin, SnxA-GFP, SnxA-mCherry, and *fab1-ha* containing yeast were obtained by genome integration of corresponding plasmids. Plasmids used in this study are listed in Table S1.

SnxA containing plasmids were generated as follows: The SnxA-GFP sequence (Vines et al., 2023) was codon optimized for yeast and synthesized in the pUC18 plasmid under XbaI and SalI restriction sites (pHM001). SnxA-GFP fragment excised by XbaI and SalI was subsequently ligated under TEF1 promoter and CYC1 terminator sequences on p415TEF. The TEF1-SnxA-GFP-CYC fragment was then excised using KpnI and SacI and subcloned into pRS306 to yield pHM007. pHM007 is linearized by ApaI digestion for integration into the *URA3* locus. To construct the SnxA-mCherry containing plasmid, mCherry excised from pAK011 using XhoI and BamHI was used to replace GFP in pHM001. SnxA-mCherry fragment was subcloned under TEF1 promoter and CYC1 terminator sequences on p415TEF using XbaI and SalI. Finally, the TEF1-SnxA-mCherry-CYC fragment was excised using KpnI and SacI and ligated into pRS306 and pRS304 to yield pHM009 and pHM011 respectively. pHM009 was digested with ApaI to integrate into *URA3* locus, whereas pHM011 was digested with Kpn2I for integration into *TRP1* locus.

For integration of Atg18 mutants into the endogenous locus: ∼4,3kb Atg18-eGFP-klTRP1 fragment amplified from the chromosomal DNA of Atg18-eGFP-klTRP1 containing yeast strain was cloned into pRS315/XmaI&SacII to obtain pCD001. FGGG and Sloop variant containing plasmids (pCD002, pCD003) were obtained through site-directed mutagenesis of pCD001. Atg18-eGFP-klTRP1 fragments of these plasmids were amplified and used as cassettes to replace *atg18Δ::hphNT1* in CDY53.

### Fluorescence Imaging

All microscopy experiments were conducted using a Carl Zeiss Axio Observer 7 motorized inverted epifluorescence microscope, equipped with a Colibri 7 LED light source, Axiocam 702 Monochrome camera, 63× Plan Apochromat immersion oil objective lens, Zeiss filter sets 95 (GFP/Cherry) and 44 (FITC), and an XL incubation and climate chamber.

For time-lapse experiments, cells were grown in filter-sterilized SC-complete media to the log phase and attached to glass-bottom Petri dishes (WVR 10810-054 Matsunami) as previously described (Caydasi and Pereira, 2017). Briefly, the center of the glass dish was coated with 6% Concanavalin A (*Canavalia ensiformis* Jack Bean, type IV Sigma C2010-G) for 10 minutes, then rinsed with sterile water. Next, 200 µL of a logarithmic phase culture was added to the center of the dish and incubated at 30°C for 30 minutes. Non-attached cells were removed by washing with pre-warmed media. The dish was filled with pre-warmed media, placed on the microscope stage, and allowed to sit for 1 hour before starting the experiment. The microscope chamber was preheated to 30°C, 2-3 hours before the experiment. For time-lapse experiments that monitored response to hyperosmotic shock, the glass-bottom petri dish was initially filled halfway with prewarmed media. Once the experiment commenced, 1.8M NaCl dissolved in filter-sterilized SC-Complete media was carefully added at 1:1 ratio (v:v) into the dish on the microscope.

Time-lapse experiments involving SnxA-GFP without hyperosmotic shock, 3xmCherry-Tubulin, and v-SEP-mCherry *atg18*Δ were conducted with images taken at 3-minute intervals. For Atg18-mNG Vph1-3xmCherry, images were captured at 5-minute intervals. Imaging frequency for GFP-Tubulin, SnxA-GFP with hyperosmotic shock and v-SEP-mCherry WT, the interval was set at 1-minute intervals.

Still images of cells were captured without cell fixation. Briefly, 1 mL of log-phase culture grown in filter sterilized SC-complete was centrifuged at 3200 rpm for 2 minutes, and cells were resuspended in an appropriate volume of SC-Complete medium before wet mount preparation. A maximum of 3 images were acquired immediately from each slide to prevent stressing of the cells and photobleaching of the signals. For the still images involving hyperosmotic shock, cell suspensions were mixed with 1.8M NaCl containing filter sterilized SC-Complete media at a ratio of 1:1 (v:v) and imaged between 5-10 minutes after salt addition.

Imaging conditions were as follows: v-SEP-mCherry was imaged using the FITC channel at 5% and the mCherry channel at 10%. SnxA-GFP was captured using the FITC channel at 10%. Vph1-3xmCherry and 3xmCherry-Tubulin were imaged using the mCherry channel at 10%. SnxA-mCherry utilized the mCherry channel at 20%, GFP-Tubulin used the FITC channel at 20%, and Atg18-mNG was imaged using FITC at 15%. 2×2 binning was applied during acquisition. For still images, 13 z-stacks of 0,3 µm thickness were acquired per image. For time-lapse experiments, 9 z-stacks of 0,4 µm thickness were acquired per image.

### Assessing timing of SnxA and Atg18 vacuole localization

The temporal localization of SnxA and Atg18 on vacuole membranes with respect to vacuole segregation and spindle elongation was judged by eye through monitoring of individual cells from the time-lapse series using Image J (NIH) software.

For the timing with respect to vacuole segregation, all cells in which segregation of the vacuole can be observed were inspected. Vacuole segregation process usually involves vesicle movement from the mother to daughter at several steps. The end of vacuole segregation was marked as the point after which vesicle movement was not observed. Timepoints at which a clear SnxA or Atg18 fluorescent signal was detected at the mother or daughter vacuole peripheries were noted. The time points of signal appearance and/or disappearance were recorded. Signal persistence was calculated as the duration between signal appearance and disappearance. Timing of signal appearance was calculated as the time between end of vacuole segregation and signal appearance.

For the timing with respect to spindle elongation, all cells that underwent both the metaphase-to-anaphase transition and the spindle breakdown during the time-lapse duration were analyzed. The timepoints for anaphase onset and spindle breakdown were determined by observing when rapid spindle elongation and spindle breakdown first occurred. The timing of SnxA-GFP and Atg18-mNG signal appearance on the daughter vacuole was calculated with respect to the timing of anaphase onset (t=0). The timing of SnxA-GFP and Atg18-mNG signal disappearance from the daughter vacuole was calculated with respect to the timing of spindle breakdown. To be able to show both timings on the same graph, spindle breakdown timing was normalized to 20 minutes for each cell.

### Assessing vacuole periphery localization via EzColocalization

To calculate Pearson Correlation Coefficient (PCC) values of Atg18/SnxA/FYVE and Vph1 signals, z-projection were performed by summing 4 z-stacks, in which the vacuole was in focus based on the Vph1 reference. Using two reporter inputs and the ROI manager for cell identification, the EzColocalization plugin ImageJ (NIH, Bethesda, Maryland, United States) was used to determine the PCC between Atg18/SnxA/FYVE and Vph1. First, the vacuoles were manually selected from the Vph1 channel using the oval selection tool and added to the ROI manager. In the EzColocalization window, PCC was selected, with the ROI set for cell identification.

PCC values from time-lapse experiments were obtained in the same way, this time the process was repeated per cell for multiple consequent time points. The time point of end of vacuole segregation (see the section above) was also noted based on Vph1 signal. The PCC values were then aligned according to this timing, mean PCC per timepoint were plotted.

### Measurement of vacuole diameter

Vacuole diameters were measured in ImageJ, by drawing a line across the vacuole based on the vacuole marker Vph1 on maximum projected images and then using the measure tool.

### Assessing Protein Asymmetry on Mother and Daughter Vacuoles

For analysis of protein asymmetry on mother and daughter vacuoles, z-stacks were sum-projected. Daughter to mother ratios of proteins of interest (Figure 4B) were determined using the following 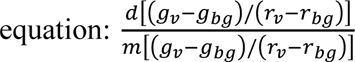; where *m* and *d* represent mother and daughter vacuoles respectively, *g* denotes mean fluorescence intensity of mNG/GFP (green signal), *r* means mean fluorescence intensity of mCherry (red signal), *v* indicates signal on the vacuole, and *gb* and *rb* represents the cytosolic signal (background) in the green and red channels respectively. Vacuoles were selected using oval selection tool based on Vph1-3xmCherry signals. Briefly, the same area was used to measure the mean fluorescence intensity of Vph1-3xmCherry and the protein of interest tagged with GFP/mNG. Mean fluorescence intensities were corrected for the background signal by subtracting the cytoplasmic signal from the vacuolar signal. Signal of the protein of interest was normalized by the Vph1-3xmCherry mean fluorescence intensity to obtain the relative levels of protein of interest per unit vacuole membrane (Vph1-3xmCherry serves as a reference). Normalized values of daughter and mother vacuoles were divided to obtain daughter to mother ratios of the protein per unit vacuole membrane.

### Analysis of vacuolar pH

For analysis of vacuolar pH, z-stacks were sum-projected using ImageJ. Using the oval selection tool, the central region of the vacuole was selected based on mCherry signal. Mean fluorescence intensity of this area was measured in both the v-SEP and mCherry channels. The mean fluorescence intensity of an area free from cells was also measured as the background. Mean fluorescence intensities of v-SEP and mCherry were corrected for background signal by subtracting the background signal from the vacuolar signal. vSEP/mCherry was calculated by dividing the corrected v-SEP signal by the corrected mCherry signal. vSEP/mCherry ratios from time-lapse experiments were obtained in the same way. The time point of end of vacuole segregation was also noted based on mCherry signal. vSEP/mCherry ratios were then aligned according to this time point, vSEP/mCherry ratios per timepoint were plotted.

### pH sensitivity and calibration curves for vSEP, mCherry and vSEP/mCherry

Cells for pH sensitivity and calibration experiments were prepared as previously described with minor modifications (Okreglak et al., 2023). Briefly, MHY283 strain that has *TEFprom-vSEP-mCherry-Cyc1term* integrated into its HO-locus was grown in 10 ml SC-complete medium to log phase. Cells were pelleted at 3200 rpm for 2 min and resuspended in 500 µl dH_2_O. 10 µl of cell suspension was pipetted in 90 µl Carmody buffers (Carmody, 1961) at different pH (3- to-10). To this mixture 1 µl of 10% digitonin (in DMSO) was added, mixed and incubated for 20 min at room temperature. Using the cell suspensions, wet mounts were prepared for microscopy. Images were acquired at the brightfield, vSEP and mCherry channels. Signals in the vacuole lumen were quantified as described above (see analysis of vacuolar pH). Corrected vSEP and mCherry mean signal intensities of the vacuole lumen were either plotted individually, or vSEP/mCherry ratios were graphed. Lines (pH 5-to-8 range) were fitted with simple linear regression model in Prism.

### Protein methods

The total yeast cell protein precipitation, sample preparation, and western blotting were performed as previously described (Meitinger et al., 2016). Primary antibodies used were rabbit anti-GFP (Abcam, ab290) and rabbit anti-Tubulin (Abcam, EPR13799). The secondary antibody was goat anti-rabbit HRP-conjugated (Advansta #R-05072-500). Chemiluminescence signals were detected using the Bio-Rad Chemidoc MP system. GFP and tubulin band intensities were measured using ImageJ and corrected for membrane background signal. The area sizes measured were kept constant across all time points. Relative GFP levels were calculated by dividing the background-corrected GFP band intensities by the corresponding background-corrected tubulin band intensities.

### Spot assay for cell growth comparison on agar plates

Yeast cultures were grown in rich media at 30°C until reaching the stationary phase. The cultures were then adjusted to an OD_600_ of 1, and 10-fold serial dilutions were prepared using sterile 1xPBS. Ten microliters of each serial dilution were spotted onto YPD agar plates and incubated at appropriate temperatures for 1-3 days.

### Data presentation and Statistical analysis

Images were visualized and analyzed using ImageJ, figures assembled in Photoshop, and labeled in Illustrator (Adobe, San Jose, CA, USA). Graphs were plotted using Prism 10.2.3 (GraphPad, Le Jolla, CA, USA), and statistical significance was determined using one-way ANOVA or unpaired two-tailed t-tests as indicated in the figure legends. In Box-and-whisker plots, boxes extend from the 25th to 75th percentiles, whiskers point to the 10^th^ and 90^th^ percentiles, and lines inside the boxes are the median.

## Acknowledgements

This study was supported by EMBO installation Grant No. 3918, and TÜBITAK Grant No. 219Z100. MH and MK were funded by TÜBITAK Grant No. 122Z887, 219Z100 and 118Z979. We would like to thank to Gislene Pereira (University of Heidelberg, Germany), Elmar Schiebel (University of Heidelberg, Germany), Andreas Mayer (University of Lausanne, Switzerland), Keisuke Obara (Nagoya University, Japan), Yoshinori Ohsumi (Tokyo Institute of Technology, Japan), Daniel Gottschling (Calico Life Sciences, USA) and Voytek Okreglak (Altos Labs, USA) for sharing yeast strains and plasmids. We are grateful to Jason King (University of Sheffield, UK) for sharing the sequence of the SnxA reporter prior to its release.

## Author Contributions

MH, MK and CD designed and performed the experiments. AKC acquired funding, designed experiments, supervised MH, MK and CD. All authors contributed to the writing of the manuscript.

---

**Figure S1- Properties of SnxA-GFP and Atg18-mNG as in vivo PI(3,5)P_2_ sensors**

A. SnxA-GFP expression levels in different strains. A representative immunoblot was shown in the bottom. Tub1 was used as a loading control. The graph on the top of each band shows the average of anti-GFP to anti-Tub1 band intensity ratios of from 4 experiments. Error bars: standard error of the mean. Strains used: MHY286, MHY288, MHY293, MHY304, MHY305, MHY303, MHY302, and AAY007.

B. Box-and-whisker graph and representative images of vacuole diameters in wildtype (WT), *vac7*Δ and SnxA-GFP containing yeast cells (BBY070, MHY286, and SEY256). Data shown comes from 3 independent experiments. Boxes show 25th and 75th percentiles, whiskers indicate 10^th^ and 90^th^ percentiles, and the line in the boxes represents the median. Ordinary one-way ANOVA was performed with Fisher’s LSD test. **** p<0.0001, ns: non-significant with p=0.7079. Scale bars: 3 μm. BF: bright field.

C. Spot assays that show growth of cells with (+SnxA-GFP) or without SnxA-GFP. Serial dilutions of indicated strains (ESM356-1 and MHY286) were spotted on YPD plates and incubated at indicated temperatures.

D-F. Vacuolar localization patterns of SnxA.

G. Analysis Atg18-mNG colocalization with Vph1-3xmCherry upon hyperosmotic shock. Atg18, Atg8-Sloop mutant, and Atg18-FGGG mutant C-terminally fused with mNeongreen were analyzed (MUK089, MUK090, and MUK091). Atg18-FGGG mutant served as a negative control for vacuole periphery localization. The PCC value of 0.5, indicated with a red dashed line, was selected as a threshold for vacuole membrane localization based on the PCC data of Atg18-FGGG. Data comes from 2 independent experiments.

H. Atg18-mNG vacuole localization reflects PI(3,5)P_2_ levels. PI(3,5)P_2_ levels were manipulated through the use of mutants that are deficient in PI(3,5)P_2_ (*vac7*Δ and *vac14*Δ), as well as through the use of *fab1-ha* and 0.9M NaCl treatment in combination with wildtype and *vac7*Δ and *vac14*Δ mutants. PCC values of mNG and Vph1-3xmCherry colocalization were plotted, and representative images were also shown. Analyzed strains: MUK089, MUK098, MUK081, MUK113, MUK080, and MUK110. Data comes from 3 independent experiments. In box-and-whiskers graphs boxes show the 25th and 75th percentiles, whiskers point to the 10^th^ and 90^th^ percentile, and the line in the boxes represents the median. In G and H, ordinary one-way ANOVA was performed with Fisher’s LSD test. **** p<0.0001, and ns: non-significant with p=0.9908. Scale bars are 3 μm.

**Figure S2. pH calibration curves, and the effect of carbon source, amino acids and Pho85 on SnxA localization.**

A-C. pH sensitivity of vSEP (A), mCherry (B) and pH calibration curve for vSEP/mCherry (C) Pink shaded areas are where vSEP intensity and vSEP/mCherry ratios linearly corralates with pH. Green shaded area in C marks the vSEP/mCherry ratios within the linear range. Data comes from 2 independent experiments. In each experiment, at least 100 vacuoles were quantified per different pH. Error bars are standard deviation. Fitted lines were obtained with simple linear regression model in Prism.

D-F. Analysis of SnxA-GFP localization at the vacuole periphery with respect to the carbon source (D), amino acid supplementation ( E) and presence of Pho85 (F). Representative still images were shown and PCC values were ploted. Data comes from 3 independent experiments. In box-and-whiskers graphs boxes show the 25th and 75th percentiles, whiskers point to the 10^th^ and 90^th^ percentile, and the line in the boxes represents the median. Ordinary one-way ANOVA was performed with Fisher’s LSD test. **** p<0.0001. Scale bars are 3 μm.

## Supplementary Table

Table S1. List of strains and plasmids used in the study

## Description of Supplementary movies

**Supplementary movie 1:** Timelapse movie of SnxA-GFP Vph1-3xmCherry containing yeast strain (MHY286) upon 0.9M NaCl treatment between the first and second timepoints. The channels from left to right are GFP, mCherry, and merge.

**Supplementary movie 2:** Timelapse movie of SnxA-GFP Vph1-3xmCherry containing yeast strain (MHY286). The channels from left to right are GFP, mCherry, and merge.

**Supplementary movie 3:** Timelapse movie of SnxA-mCherry GFP-Tub1 containing yeast strain (MHY295). The channels from left to right are mCherry, GFP and merge.

**Supplementary movie 4:** Timelapse movie of Atg18-mNeongreen containing yeast strain (MUK089). The channels from left to right are mNeongreen, mCherry, and merge.

**Supplementary movie 5:** Timelapse movie of vSEP-mCherry containing yeast strain (MHY283). The channels from left to right are vSEP, mCherry, and merge.

**Supplementary movie 6:** Timelapse movie of vSEP-mCherry containing *atg18*Δ yeast strain (MHY277). The channels from left to right are vSEP, mCherry, and merge.

## References

Banerjee, S., K. Clapp, M. Tarsio, and P.M. Kane. 2019. Interaction of the late endo-lysosomal lipid PI(3,5)P2 with the Vph1 isoform of yeast V-ATPase increases its activity and cellular stress tolerance. J Biol Chem. 294:9161–9171.

Barlow-Busch, I., A.L. Shaw, and J.E. Burke. 2023. PI4KA and PIKfyve: Essential phosphoinositide signaling enzymes involved in myriad human diseases. Curr Opin Cell Biol. 83:102207.

Botman, D., D.H. de Groot, P. Schmidt, J. Goedhart, and B. Teusink. 2019. In vivo characterisation of fluorescent proteins in budding yeast. Sci Rep. 9:2234.

Bridges, D., J.T. Ma, S. Park, K. Inoki, L.S. Weisman, and A.R. Saltiel. 2012. Phosphatidylinositol 3,5-bisphosphate plays a role in the activation and subcellular localization of mechanistic target of rapamycin 1. Mol Biol Cell. 23:2955–2962.

Busse, R.A., A. Scacioc, R. Krick, A. Perez-Lara, M. Thumm, and K. Kuhnel. 2015. Characterization of PROPPIN-Phosphoinositide Binding and Role of Loop 6CD in PROPPIN-Membrane Binding. Biophys J. 108:2223–2234.

Carmody, W.R. 1961. Easily prepared wide range buffer series. Journal of Chemical Education 38.

Caydasi, A.K., A. Khmelinskii, R. Duenas-Sanchez, B. Kurtulmus, M. Knop, and G. Pereira. 2017. Temporal and compartment-specific signals coordinate mitotic exit with spindle position. Nat Commun. 8:14129.

Caydasi, A.K., and G. Pereira. 2017. Evaluation of the Dynamicity of Mitotic Exit Network and Spindle Position Checkpoint Components on Spindle Pole Bodies by Fluorescence Recovery After Photobleaching (FRAP). Methods Mol Biol. 1505:167–182.

Chen, Z., P.C. Malia, R. Hatakeyama, R. Nicastro, Z. Hu, M.P. Peli-Gulli, J. Gao, T. Nishimura, E. Eskes, C.J. Stefan, J. Winderickx, J. Dengjel, C. De Virgilio, and C. Ungermann. 2021. TORC1 Determines Fab1 Lipid Kinase Function at Signaling Endosomes and Vacuoles. Curr Biol. 31:297–309 e298.

Dove, S.K., F.T. Cooke, M.R. Douglas, L.G. Sayers, P.J. Parker, and R.H. Michell. 1997. Osmotic stress activates phosphatidylinositol-3,5-bisphosphate synthesis. Nature. 390:187–192.

Dove, S.K., R.K. McEwen, A. Mayes, D.C. Hughes, J.D. Beggs, and R.H. Michell. 2002. Vac14 controls PtdIns(3,5)P(2) synthesis and Fab1-dependent protein trafficking to the multivesicular body. Curr Biol. 12:885–893.

Dove, S.K., R.C. Piper, R.K. McEwen, J.W. Yu, M.C. King, D.C. Hughes, J. Thuring, A.B. Holmes, F.T. Cooke, R.H. Michell, P.J. Parker, and M.A. Lemmon. 2004. Svp1p defines a family of phosphatidylinositol 3,5-bisphosphate effectors. EMBO J. 23:1922–1933.

Duex, J.E., J.J. Nau, E.J. Kauffman, and L.S. Weisman. 2006a. Phosphoinositide 5-phosphatase Fig 4p is required for both acute rise and subsequent fall in stress-induced phosphatidylinositol 3,5-bisphosphate levels. Eukaryotic cell. 5:723–731.

Duex, J.E., F. Tang, and L.S. Weisman. 2006b. The Vac14p-Fig4p complex acts independently of Vac7p and couples PI3,5P2 synthesis and turnover. J Cell Biol. 172:693–704.

Efe, J.A., R.J. Botelho, and S.D. Emr. 2007. Atg18 regulates organelle morphology and Fab1 kinase activity independent of its membrane recruitment by phosphatidylinositol 3,5-bisphosphate. Mol Biol Cell. 18:4232–4244.

Finnigan, G.C., G.E. Cronan, H.J. Park, S. Srinivasan, F.A. Quiocho, and T.H. Stevens. 2012. Sorting of the yeast vacuolar-type, proton-translocating ATPase enzyme complex (V-ATPase): identification of a necessary and sufficient Golgi/endosomal retention signal in Stv1p. J Biol Chem. 287:19487–19500.

Gary, J.D., T.K. Sato, C.J. Stefan, C.J. Bonangelino, L.S. Weisman, and S.D. Emr. 2002. Regulation of Fab1 phosphatidylinositol 3-phosphate 5-kinase pathway by Vac7 protein and Fig4, a polyphosphoinositide phosphatase family member. Mol Biol Cell. 13:1238–1251.

Gary, J.D., A.E. Wurmser, C.J. Bonangelino, L.S. Weisman, and S.D. Emr. 1998. Fab1p is essential for PtdIns(3)P 5-kinase activity and the maintenance of vacuolar size and membrane homeostasis. J Cell Biol. 143:65–79.

Gillooly, D.J., I.C. Morrow, M. Lindsay, R. Gould, N.J. Bryant, J.M. Gaullier, R.G. Parton, and H. Stenmark. 2000. Localization of phosphatidylinositol 3-phosphate in yeast and mammalian cells. EMBO J. 19:4577–4588.

Giridharan, S.S.P., G. Luo, P. Rivero-Rios, N. Steinfeld, H. Tronchere, A. Singla, E. Burstein, D.D. Billadeau, M.A. Sutton, and L.S. Weisman. 2022. Lipid kinases VPS34 and PIKfyve coordinate a phosphoinositide cascade to regulate retriever-mediated recycling on endosomes. Elife. 11.

Gopaldass, N., B. Fauvet, H. Lashuel, A. Roux, and A. Mayer. 2017. Membrane scission driven by the PROPPIN Atg18. EMBO J. 36:3274–3291.

Han, B.K., and S.D. Emr. 2011. Phosphoinositide [PI(3,5)P2] lipid-dependent regulation of the general transcriptional regulator Tup1. Genes Dev. 25:984–995.

Harding, T.M., A. Hefner-Gravink, M. Thumm, and D.J. Klionsky. 1996. Genetic and phenotypic overlap between autophagy and the cytoplasm to vacuole protein targeting pathway. J Biol Chem. 271:17621–17624.

Henderson, K.A., A.L. Hughes, and D.E. Gottschling. 2014. Mother-daughter asymmetry of pH underlies aging and rejuvenation in yeast. Elife. 3:e03504.

Huda, M., S.N. Bektas, B. Bekdas, and A.K. Caydasi. 2023. The signalling lipid PI3,5P(2) is essential for timely mitotic exit. Open Biol. 13:230125.

Hughes, A.L., and D.E. Gottschling. 2012. An early age increase in vacuolar pH limits mitochondrial function and lifespan in yeast. Nature. 492:261–265.

Janke, C., M.M. Magiera, N. Rathfelder, C. Taxis, S. Reber, H. Maekawa, A. Moreno-Borchart, G. Doenges, E. Schwob, E. Schiebel, and M. Knop. 2004. A versatile toolbox for PCR-based tagging of yeast genes: new fluorescent proteins, more markers and promoter substitution cassettes. Yeast. 21:947–962.

Jin, N., K. Mao, Y. Jin, G. Tevzadze, E.J. Kauffman, S. Park, D. Bridges, R. Loewith, A.R. Saltiel, D.J. Klionsky, and L.S. Weisman. 2014. Roles for PI(3,5)P2 in nutrient sensing through TORC1. Mol Biol Cell. 25:1171–1185.

Jin, Y., N. Jin, Y. Oikawa, R. Benyair, M. Koizumi, T.E. Wilson, Y. Ohsumi, and L.S. Weisman. 2022. Bur1 functions with TORC1 for vacuole-mediated cell cycle progression. EMBO Rep. 23:e53477.

Jin, Y., and L.S. Weisman. 2015. The vacuole/lysosome is required for cell-cycle progression. Elife. 4.

Jones, D.R., A. Gonzalez-Garcia, E. Diez, A.C. Martinez, A.C. Carrera, and I. Merida. 1999. The identification of phosphatidylinositol 3,5-bisphosphate in T-lymphocytes and its regulation by interleukin-2. J Biol Chem. 274:18407–18413.

Kane, P.M. 1995. Disassembly and reassembly of the yeast vacuolar H(+)-ATPase in vivo. J Biol Chem. 270:17025–17032.

Kane, P.M. 2016. Proton Transport and pH Control in Fungi. Adv Exp Med Biol. 892:33–68.

Knop, M., K. Siegers, G. Pereira, W. Zachariae, B. Winsor, K. Nasmyth, and E. Schiebel. 1999. Epitope tagging of yeast genes using a PCR-based strategy: more tags and improved practical routines. Yeast. 15:963–972.

Lees, J.A., P. Li, N. Kumar, L.S. Weisman, and K.M. Reinisch. 2020. Insights into Lysosomal PI(3,5)P(2) Homeostasis from a Structural-Biochemical Analysis of the PIKfyve Lipid Kinase Complex. Mol Cell. 80:736–743 e734.

Li, S.C., T.T. Diakov, T. Xu, M. Tarsio, W. Zhu, S. Couoh-Cardel, L.S. Weisman, and P.M. Kane. 2014. The signaling lipid PI(3,5)P(2) stabilizes V(1)-V(o) sector interactions and activates the V-ATPase. Mol Biol Cell. 25:1251–1262.

McCartney, A.J., Y. Zhang, and L.S. Weisman. 2014a. Phosphatidylinositol 3,5-bisphosphate: low abundance, high significance. BioEssays : news and reviews in molecular, cellular and developmental biology. 36:52–64.

McCartney, A.J., S.N. Zolov, E.J. Kauffman, Y. Zhang, B.S. Strunk, L.S. Weisman, and M.A. Sutton. 2014b. Activity-dependent PI(3,5)P2 synthesis controls AMPA receptor trafficking during synaptic depression. Proc Natl Acad Sci U S A. 111:E4896–4905.

Meitinger, F., S. Palani, and G. Pereira. 2016. Detection of Phosphorylation Status of Cytokinetic Components. Springer New York. 219–237.

Obara, K., T. Noda, K. Niimi, and Y. Ohsumi. 2008. Transport of phosphatidylinositol 3-phosphate into the vacuole via autophagic membranes in Saccharomyces cerevisiae. Genes Cells. 13:537–547.

Okreglak, V., R. Ling, M. Ingaramo, N.H. Thayer, A. Millett-Sikking, and D.E. Gottschling. 2023. Cell cycle-linked vacuolar pH dynamics regulate amino acid homeostasis and cell growth. Nat Metab. 5:1803–1819.

Parra, K.J., and P.M. Kane. 1998. Reversible association between the V1 and V0 domains of yeast vacuolar H+-ATPase is an unconventional glucose-induced effect. Mol Cell Biol. 18:7064–7074.

Ruckenstuhl, C., C. Netzberger, I. Entfellner, D. Carmona-Gutierrez, T. Kickenweiz, S. Stekovic, C. Gleixner, C. Schmid, L. Klug, A.G. Sorgo, T. Eisenberg, S. Buttner, G. Marino, R. Koziel, P. Jansen-Durr, K.U. Frohlich, G. Kroemer, and F. Madeo. 2014. Lifespan extension by methionine restriction requires autophagy-dependent vacuolar acidification. PLoS Genet. 10:e1004347.

Rutherford, A.C., C. Traer, T. Wassmer, K. Pattni, M.V. Bujny, J.G. Carlton, H. Stenmark, and P.J. Cullen. 2006. The mammalian phosphatidylinositol 3-phosphate 5-kinase (PIKfyve) regulates endosome-to-TGN retrograde transport. J Cell Sci. 119:3944–3957.

Sbrissa, D., O.C. Ikonomov, and A. Shisheva. 1999. PIKfyve, a mammalian ortholog of yeast Fab1p lipid kinase, synthesizes 5-phosphoinositides. Effect of insulin. J Biol Chem. 274:21589–21597.

Sbrissa, D., and A. Shisheva. 2005. Acquisition of unprecedented phosphatidylinositol 3,5-bisphosphate rise in hyperosmotically stressed 3T3-L1 adipocytes, mediated by ArPIKfyve-PIKfyve pathway. J Biol Chem. 280:7883–7889.

Sherman, F. 1991. [1] Getting started with yeast. Elsevier. 3–21.

Stauffer, W., H. Sheng, and H.N. Lim. 2018. EzColocalization: An ImageJ plugin for visualizing and measuring colocalization in cells and organisms. Sci Rep. 8:15764.

Takatori, S., T. Tatematsu, J. Cheng, J. Matsumoto, T. Akano, and T. Fujimoto. 2016. Phosphatidylinositol 3,5-Bisphosphate-Rich Membrane Domains in Endosomes and Lysosomes. Traffic. 17:154–167.

Tsujita, K., T. Itoh, T. Ijuin, A. Yamamoto, A. Shisheva, J. Laporte, and T. Takenawa. 2004. Myotubularin regulates the function of the late endosome through the gram domain-phosphatidylinositol 3,5-bisphosphate interaction. J Biol Chem. 279:13817–13824.

Vines, J.H., H. Maib, C.M. Buckley, A. Gueho, Z. Zhu, T. Soldati, D.H. Murray, and J.S. King. 2023. A PI(3,5)P2 reporter reveals PIKfyve activity and dynamics on macropinosomes and phagosomes. J Cell Biol. 222.

Yorimitsu, T., and D.J. Klionsky. 2005. Atg11 links cargo to the vesicle-forming machinery in the cytoplasm to vacuole targeting pathway. Mol Biol Cell. 16:1593–1605.

