## Supplementary figures for "PI(3,5)P_2_ asymmetry during mitosis is essential for asymmetric vacuolar inheritance"

In box-and-whisker graphs, boxes show the 25th and 75th percentiles, whiskers indicate the $10^{th}$ and $90^{th}$ percentiles, and the line in the boxes represents the median. In A, C, E and F, ordinary one-way ANOVA was performed with Fisher's LSD test. **** $p<0.0001$, ** $p\leq0.0093$, *** $p=0.0004$, and ns: non-significant with $p\geq0.6580$. Scale bars: 3 µm. BF: bright field.

### Figure 1

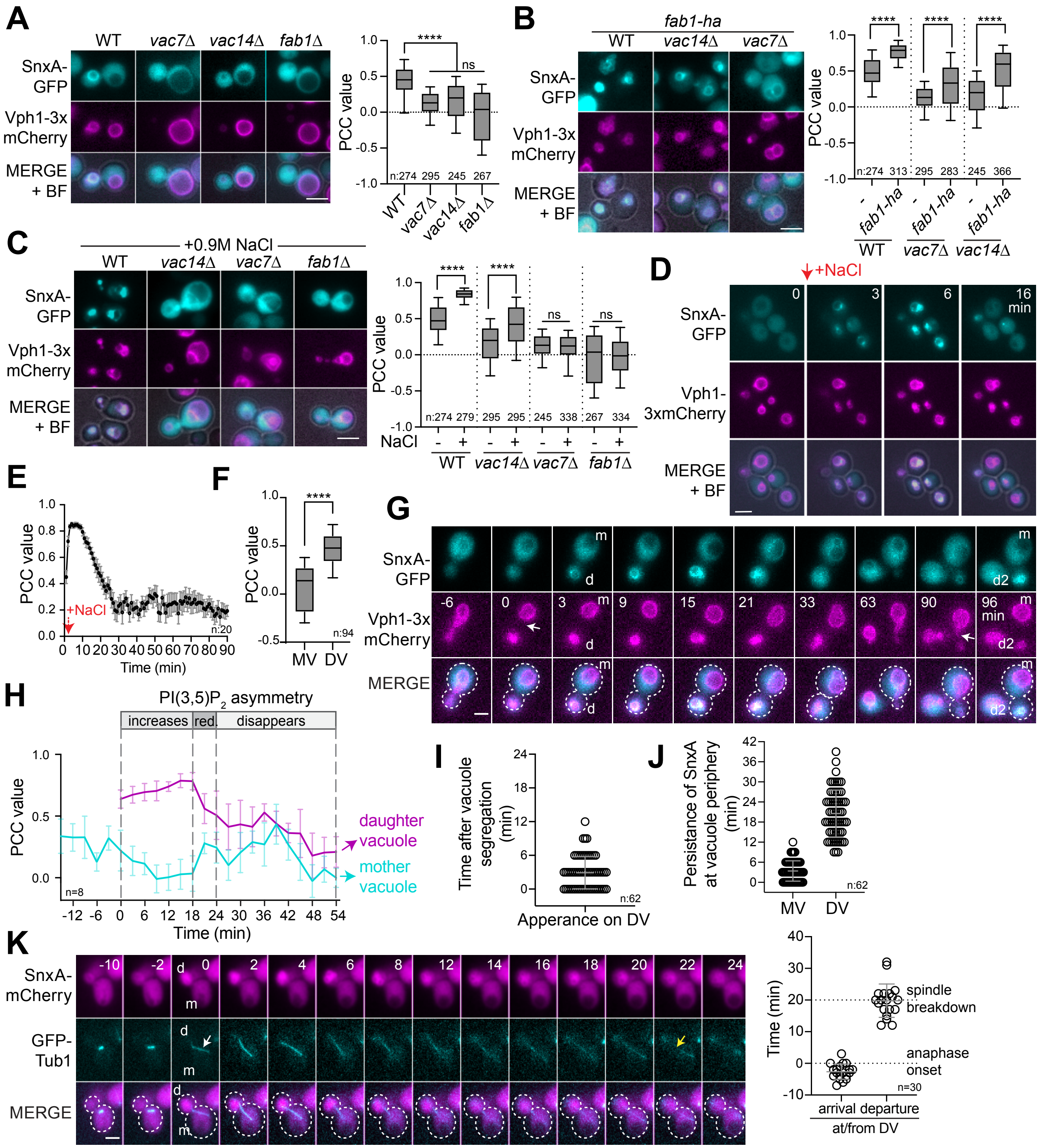

A
WT
vac7Δ
vac14Δ
fab1Δ
SnxA-GFP
Vph1-3x mCherry
MERGE + BF
PCC value
n:274 295 245 267
****
ns
B
fab1-ha
WT
vac14Δ
vac7Δ
SnxA-GFP
Vph1-3x mCherry
MERGE + BF
n:274 313 295 283 245 366
-
fab1-ha
C
+0.9M NaCl
WT
vac14Δ
vac7Δ
fab1Δ
SnxA-GFP
Vph1-3x mCherry
MERGE + BF
PCC value
n:274 279 295 295 245 338 267 334
NaCl
D
+NaCl
0
3
6
16 min
SnxA-GFP
Vph1-3xmCherry
MERGE + BF
E
PCC value
+NaCl
n:20
Time (min)
F
PCC value
n:94
MV
DV
G
SnxA-GFP
Vph1-3x mCherry
MERGE
-6
0
3
9
15
21
33
63
90
96 min
m
d
d2
H
PI(3,5)P$_2$ asymmetry
increases
red.
disappears
PCC value
n=8
daughter vacuole
mother vacuole
Time (min)
I
Time after vacuole segregation (min)
n:62
Apperance on DV
J
Persistance of SnxA at vacuole periphery (min)
MV
DV
n:62
K
SnxA-mCherry
GFP-Tub1
MERGE
-10
-2
0
2
4
6
8
12
14
16
18
20
22
24
d
m
Time (min)
spindle breakdown
anaphase onset
n=30
arrival
departure
at/from DV

### Figure 2

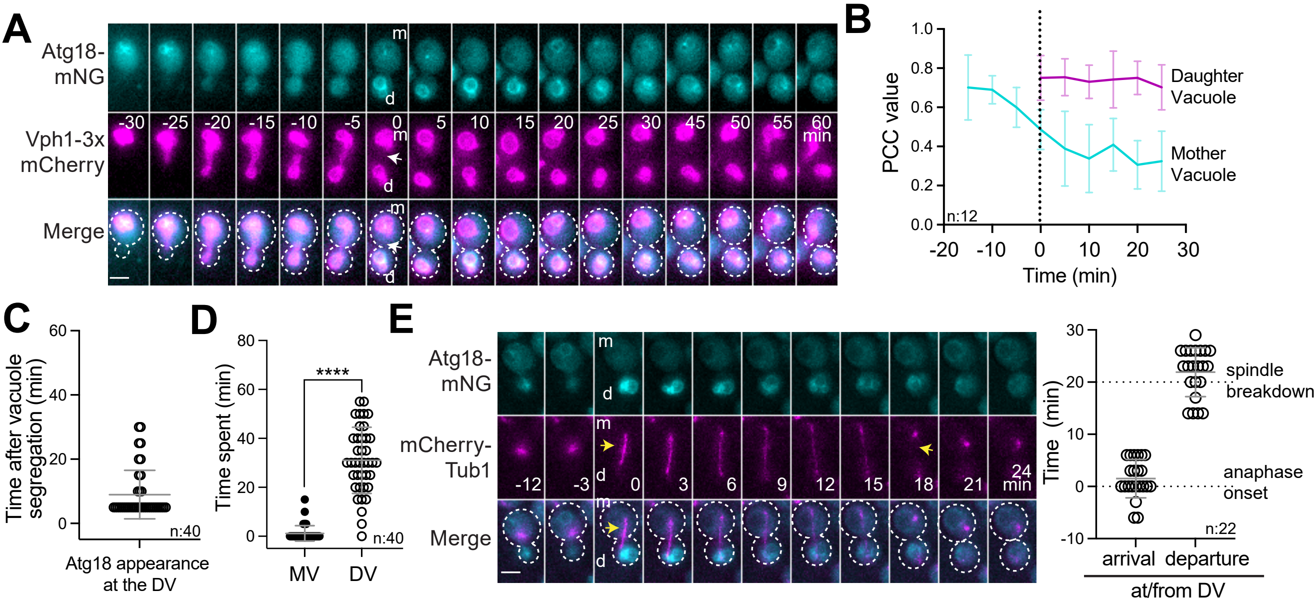
A
Atg18-mNG
Vph1-3x mCherry
Merge
m
d
-30 -25 -20 -15 -10 -5 0 5 10 15 20 25 30 45 50 55 60 min
B
PCC value
Time (min)
Daughter Vacuole
Mother Vacuole
n:12
C
Time after vacuole segregation (min)
Atg18 appearance at the DV
n:40
D
Time spent (min)
****
MV
DV
n:40
E
Atg18-mNG
mCherry-Tub1
Merge
-12 -3 0 3 6 9 12 15 18 21 24 min
Time (min)
spindle breakdown
anaphase onset
arrival
departure
at/from DV
n:22

### Figure 3

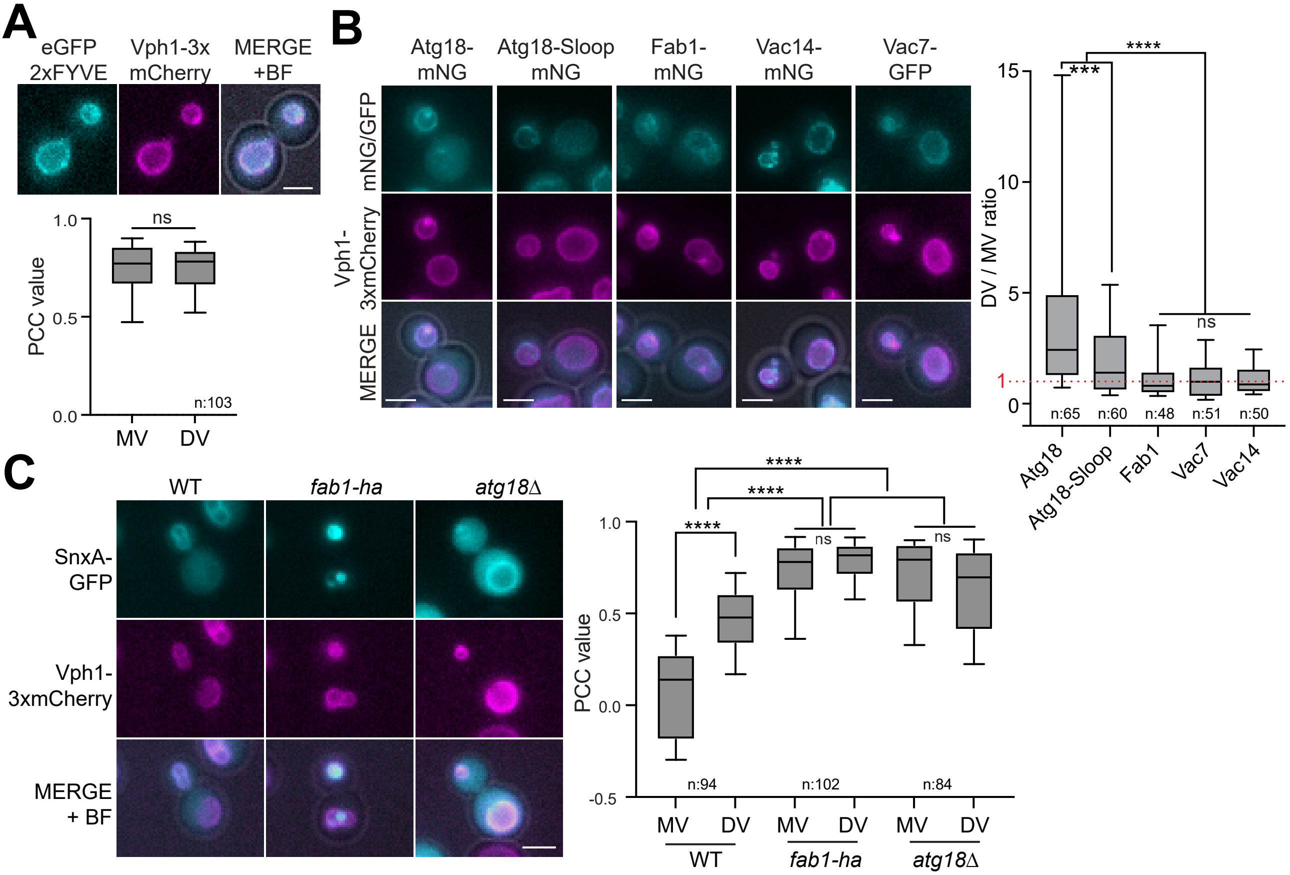
A
eGFP 2xFYVE
Vph1-3x mCherry
MERGE +BF
ns
PCC value
n:103
MV
DV
B
Atg18-mNG
Atg18-Sloop mNG
Fab1-mNG
Vac14-mNG
Vac7-GFP
mNG/GFP
Vph1-3xmCherry
MERGE
DV / MV ratio
****
***
ns
n:65
n:60
n:48
n:51
n:50
Atg18
Atg18-Sloop
Fab1
Vac7
Vac14
C
WT
fab1-ha
atg18Δ
SnxA-GFP
Vph1-3xmCherry
MERGE + BF
PCC value
****
ns
n:94
n:102
n:84
MV
DV
WT
fab1-ha
atg18Δ

### Figure 4

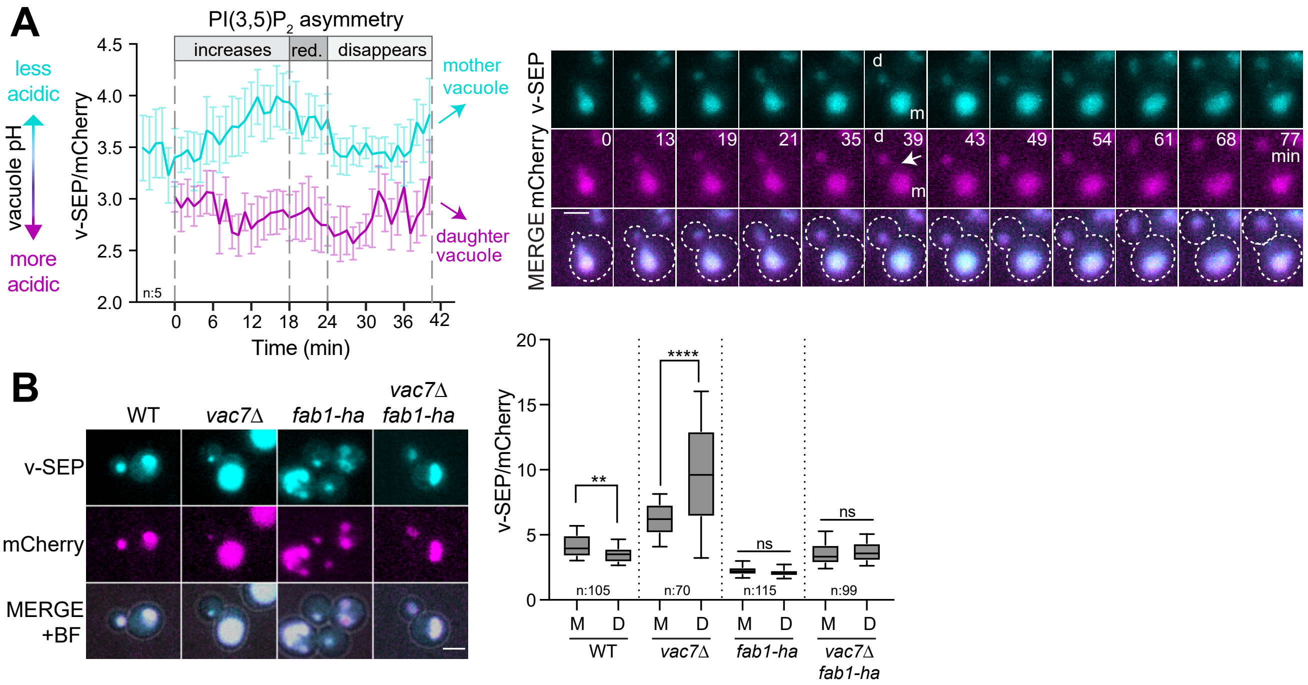

A
B
PI(3,5)P2 asymmetry
increases
red.
disappears
less acidic
vacuole pH
more acidic
v-SEP/mCherry
4.5
4.0
3.5
3.0
2.5
2.0
n:5
0
6
12
18
24
30
36
42
Time (min)
mother vacuole
daughter vacuole
v-SEP
mCherry
MERGE
d
m
13
19
21
35
39
43
49
54
61
68
77
min
WT
vac7Δ
fab1-ha
vac7Δ fab1-ha
+BF
20
15
10
5
**
****
ns
n:105
n:70
n:115
n:99
M
D

#### Figure 5

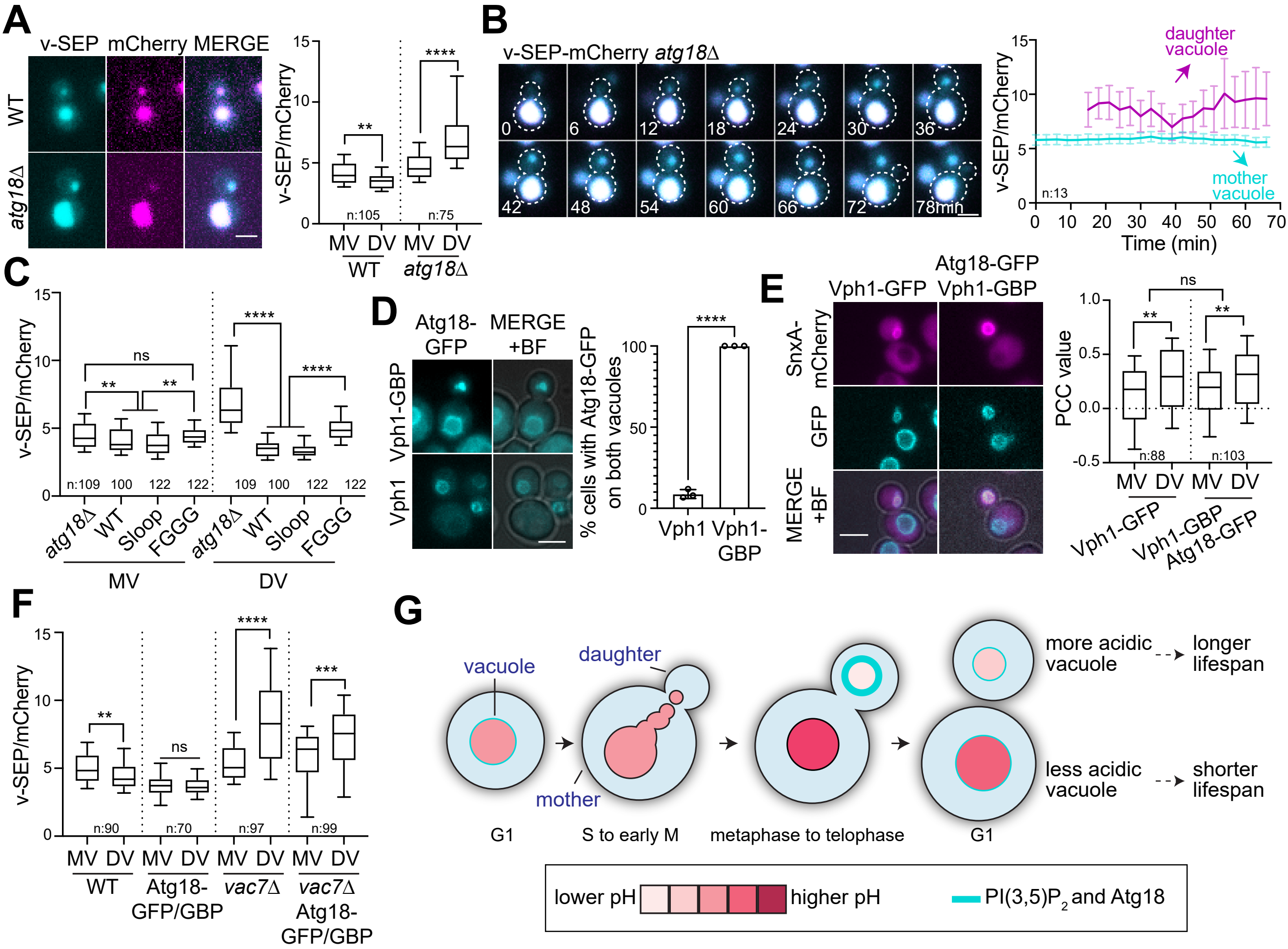

A
v-SEP mCherry MERGE
WT
atg18Δ
v-SEP/mCherry
n:105
n:75
MV DV MV DV
WT atg18Δ
B
v-SEP-mCherry atg18Δ
0 6 12 18 24 30 36
42 48 54 60 66 72 78min
daughter vacuole
mother vacuole
n:13
Time (min)
C
v-SEP/mCherry
ns
n:109 100 122 122 109 100 122 122
atg18Δ WT Sloop FGGG atg18Δ WT Sloop FGGG
MV
DV
D
Atg18-GFP
MERGE +BF
Vph1-GBP
Vph1
% cells with Atg18-GFP on both vacuoles
Vph1 Vph1-GBP
E
Vph1-GFP
Atg18-GFP Vph1-GBP
SnxA-mCherry
GFP
MERGE +BF
PCC value
ns
n:88
n:103
MV DV MV DV
Vph1-GFP
Vph1-GBP Atg18-GFP
F
v-SEP/mCherry
ns
n:90 n:70 n:97 n:99
MV DV MV DV MV DV MV DV
WT
Atg18-GFP/GBP
vac7Δ
vac7Δ Atg18-GFP/GBP
G
vacuole
daughter
mother
G1
S to early M
metaphase to telophase
G1
more acidic vacuole
longer lifespan
less acidic vacuole
shorter lifespan
lower pH
higher pH
PI(3,5)P2 and Atg18

### Figure S1

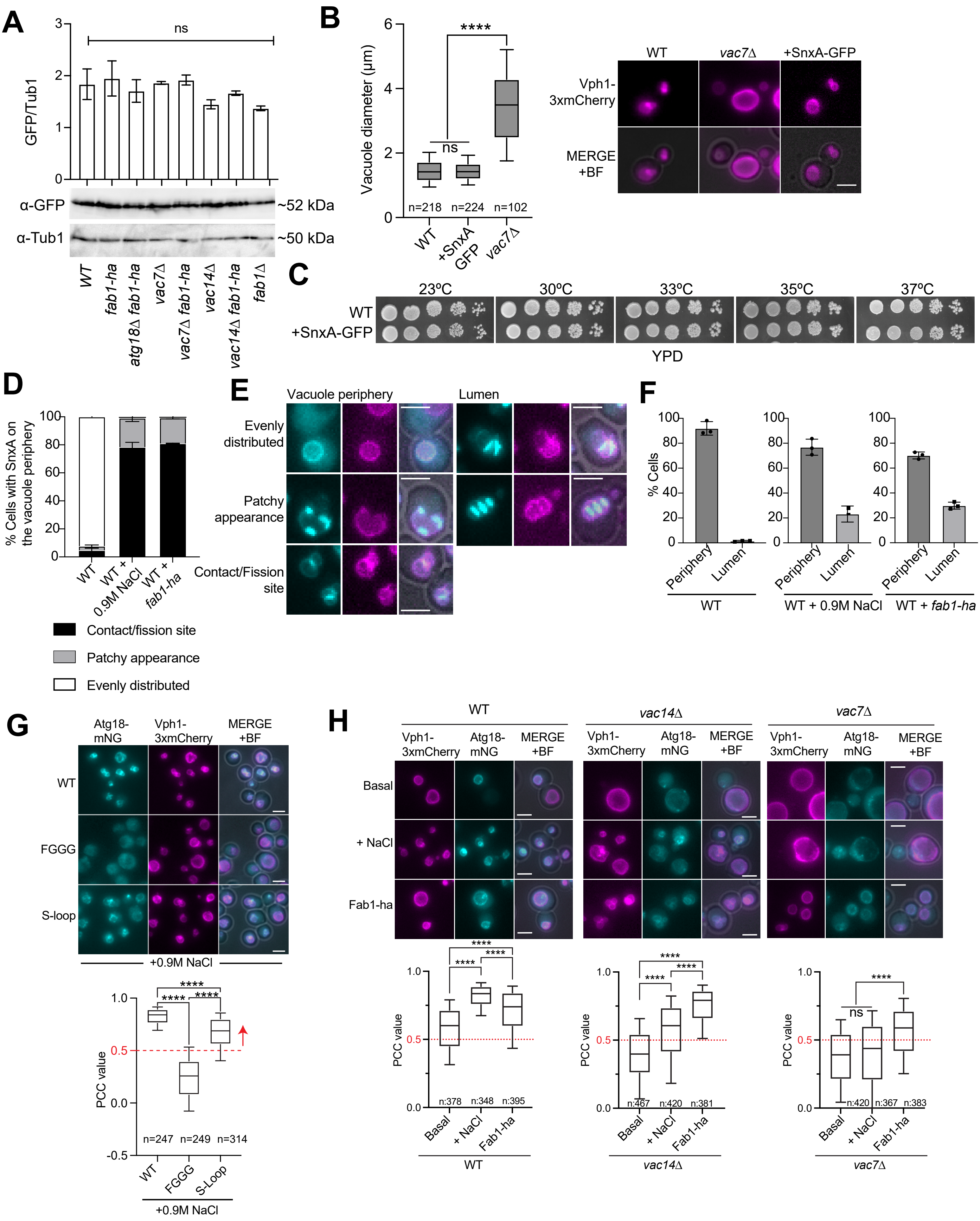
A
GFP/Tub1
ns
α-GFP
~52 kDa
α-Tub1
~50 kDa
WT
fab1-ha
atg18Δ fab1-ha
vac7Δ
vac7Δ fab1-ha
vac14Δ
vac14Δ fab1-ha
fab1Δ
B
Vacuole diameter (µm)
****
ns
n=218
n=224
n=102
WT
+SnxA GFP
vac7Δ
WT
vac7Δ
+SnxA-GFP
Vph1-3xmCherry
MERGE +BF
C
23ºC
30ºC
33ºC
35ºC
37ºC
WT
+SnxA-GFP
YPD
D
% Cells with SnxA on the vacuole periphery
WT
WT + 0.9M NaCl
WT + fab1-ha
Contact/fission site
Patchy appearance
Evenly distributed
E
Vacuole periphery
Lumen
Evenly distributed
Patchy appearance
Contact/Fission site
F
% Cells
Periphery
Lumen
WT
WT + 0.9M NaCl
WT + fab1-ha
G
Atg18-mNG
Vph1-3xmCherry
MERGE +BF
WT
FGGG
S-loop
+0.9M NaCl
PCC value
****
n=247
n=249
n=314
WT
FGGG
S-Loop
+0.9M NaCl
H
WT
vac14Δ
vac7Δ
Vph1-3xmCherry
Atg18-mNG
MERGE +BF
Basal
+ NaCl
Fab1-ha
PCC value
****
ns
n:378
n:348
n:395
n:467
n:420
n:381
n:420
n:367
n:383
Basal
+ NaCl
Fab1-ha

### Figure S2

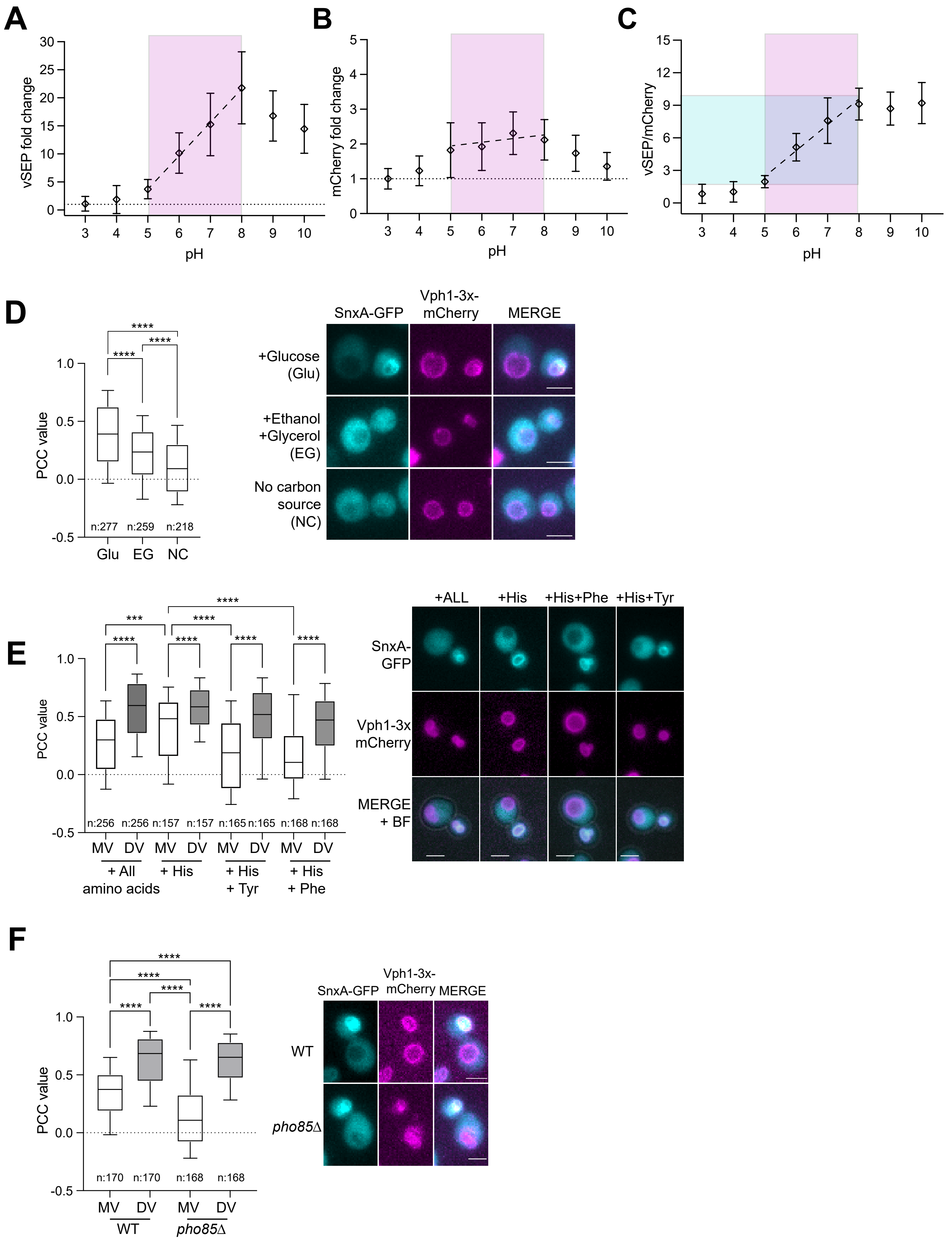

A
vSEP fold change
pH
B
mCherry fold change
pH
C
vSEP/mCherry
pH
D
PCC value
n:277
n:259
n:218
Glu
EG
NC
SnxA-GFP
Vph1-3x-mCherry
MERGE
+Glucose (Glu)
+Ethanol +Glycerol (EG)
No carbon source (NC)
E
PCC value
n:256 n:256 n:157 n:157 n:165 n:165 n:168 n:168
MV DV MV DV MV DV MV DV
+ All amino acids
+ His
+ His + Tyr
+ His + Phe
+ALL
+His
+His+Phe
+His+Tyr
SnxA-GFP
Vph1-3x mCherry
MERGE + BF
F
PCC value
n:170 n:170 n:168 n:168
MV DV MV DV
WT
pho85Δ
SnxA-GFP
Vph1-3x-mCherry
MERGE
WT
pho85Δ
