## Supplementary Table for "PI(3,5)P_2_ asymmetry during mitosis is essential for asymmetric vacuolar inheritance"

**Table S1. Yeast strains and plasmids used in this study**

| Strain Name | Genotype/description | Reference |
|---|---|---|
| ESM356 | MATa *ura3-52 leu2Δ1 his3Δ200 trp1Δ63* | Pereira et al., 2001 |
| AAY007 | MATa *ura3-52 leu2Δ1 his3Δ200 trp1Δ63 VPH1-3XmCherry-kanMX6 fab1Δ::klTRP pKL08-SnxA-GFP* | This study |
| BBY070 | MATa *ura3-52 leu2Δ1 his3Δ200 trp1Δ63 ura3-52::URA3-GFP-TUB1 VPH1-3XmCherry-kanMX6* | Huda *et al*, 2023 |
| CDY53 | MATa *ura3-52 leu2Δ1 his3Δ200 trp1Δ63 VPH1-3XmCherry-kanMX6 atg18Δ::hphNT2* | This study |
| MHY030 | MATa *ura3-52 leu2Δ1 his3Δ200 trp1Δ63 VPH1-GFP-His3MX6* leu2Δ1::LEU2-fab1-ha | This study |
| MHY223 | MATa *ura3-52 leu2Δ1 his3Δ200 trp1Δ63 ATG18-GFP-klTRP VPH1-GBP-hphNT1.* | This study |
| MHY252 | MATa *ura3-52 leu2Δ1 his3Δ200 trp1Δ63 VPH1-3XmCherry-kanMX6 pRS416-EGFP-2xFYVE* | This study |
| MHY257 | MATa *ura3-52 leu2Δ1 his3Δ200 trp1Δ63 vSEP-mCherry-kanMX6 leu2Δ1::LEU2-fab1-ha* | This study |
| MHY277 | MATa *ura3-52 leu2Δ1 his3Δ200 trp1Δ63 atg18Δ::hphNT2 vSEP-mCherry-kanMX6* | This study |
| MHY283 | MATa *ura3-52 leu2Δ1 his3Δ200 trp1Δ63 vSEP-mCherry-kanMX6* | This study |
| MHY286 | MATa *ura3-52 leu2Δ1 his3Δ200 trp1Δ63 VPH1-3XmCherry-kanMX6 ura3-52::URA3-SnxA-GFP* | This study |
| MHY288 | MATa *ura3-52 leu2Δ1 his3Δ200 trp1Δ63 VPH1-3XmCherry-kanMX6 ura3-52::URA3-SnxA-GFP leu2Δ1::LEU2-fab1-ha* | This study |
| MHY285 | MATa *ura3-52 leu2Δ1 his3Δ200 trp1Δ63 VPH1-GFP-His3MX6 ura3-52::URA3-SnxA-mCherry* | This study |
| MHY289 | MATa *ura3-52 leu2Δ1 his3Δ200 trp1Δ63 VPH1-3XmCherry-kanMX6 ura3-52::URA3-SnxA-GFP atg18Δ::hphNT2* | This study |
| MHY291 | MATa *ura3-52 leu2Δ1 his3Δ200 trp1Δ63 vSEP-mCherry-kanMX6 vac7Δ::6HAklTRP* | This study |
| MHY293 | MATa *ura3-52 leu2Δ1 his3Δ200 trp1Δ63 VPH1-3XmCherry-kanMX6 ura3-52::URA3-SnxA-GFP atg18Δ::hphNT2 leu2Δ1::LEU2-fab1-ha* | This study |
| MHY295 | MATa *ura3-52 leu2Δ1 his3Δ200 trp1Δ63 ura3-52::URA3-GFP-TUB1 trp1Δ63::TRP1-SnxA-mCherry* | This study |
| MHY296 | MATa *ura3-52 leu2Δ1 his3Δ200 trp1Δ63 vSEP-mCherry-kanMX6 vac7Δ::6HAklTRP leu2Δ1::LEU2-fab1-ha* | This study |
| MHY297 | MATa *ura3-52 leu2Δ1 his3Δ200 trp1Δ63 ATG18-GFP-klTRP VPH1-GBP-hphNT1 vSEP-mCherry-kanMX6* | This study |
| MHY298 | MATa *ura3-52 leu2Δ1 his3Δ200 trp1Δ63 ATG18-GFP-klTRP VPH1-GBP-hphNT1 ura3-52::URA3-SnxA-mCherry* | This study |
| MHY302 | MATa *ura3-52 leu2Δ1 his3Δ200 trp1Δ63 VPH1-3XmCherry-kanMX6 ura3-52::URA3-SnxA-GFP vac14Δ::6HAklTRP* | This study |
| MHY303 | MATa *ura3-52 leu2Δ1 his3Δ200 trp1Δ63 VPH1-3XmCherry-kanMX6 ura3-52::URA3-SnxA-GFP leu2Δ1::LEU2-fab1-ha vac14Δ::6HAklTRP* | This study |
| MHY304 | MATa *ura3-52 leu2Δ1 his3Δ200 trp1Δ63 VPH1-3XmCherry-kanMX6 ura3-52::URA3-SnxA-GFP vac7Δ::6HAklTRP* | This study |
| MHY305 | MATa *ura3-52 leu2Δ1 his3Δ200 trp1Δ63 VPH1-3XmCherry-kanMX6 ura3-52::URA3-SnxA-GFP leu2Δ1::LEU2-fab1-ha vac7Δ::6HAklTRP* | This study |
| MHY312 | MATa *ura3-52 leu2Δ1 his3Δ200 trp1Δ63 ATG18-GFP-klTRP VPH1-GBP-hphNT1 vac7Δ::6HAHis3MX6 vSEP-mCherry-kanMX6* | This study |
| MHY317 | MATa *ura3-52 leu2Δ1 his3Δ200 trp1Δ63 atg18Δ::hphNT2 vSEP-mCherry-kanMX6 hphNT2::ATG18-FGGG-6HAHis3MX6* | This study |
| MUK080 | MATa *ura3-52 leu2Δ1 his3Δ200 trp1Δ63 VPH1-3XmCherry-kanMX6 ATG18-mNeongreen-hphNT2 vac14Δ::6HAklTRP* | This study |
| MUK081 | MATa *ura3-52 leu2Δ1 his3Δ200 trp1Δ63 VPH1-3XmCherry-kanMX6 ATG18-mNeongreen-hphNT2 vac7Δ::6HAklTRP* | This study |
| MUK089 | MATa *ura3-52 leu2Δ1 his3Δ200 trp1Δ63 VPH1-3XmCherry-kanMX6 atg18Δ::hphNT2 ATG18-mNeongreen-hphNT2* | This study |
| MUK090 | MATa *ura3-52 leu2Δ1 his3Δ200 trp1Δ63 VPH1-3XmCherry-kanMX6 atg18Δ::hphNT2 ATG18-FGGG-mNeongreen-hphNT2* | This study |
| MUK091 | MATa *ura3-52 leu2Δ1 his3Δ200 trp1Δ63 VPH1-3XmCherry-kanMX6 atg18Δ::hphNT2 ATG18-Sloop-mNeongreen-hphNT2* | This study |
| MUK097 | MATa *ura3-52 leu2Δ1 his3Δ200 trp1Δ63 VPH1-3XmCherry-kanMX6 atg18Δ::hphNT2 ATG18-Sloop-mNeongreen-hphNT2 leu2Δ1::LEU2-fab1-* | This study |
| MUK098 | MATa *ura3-52 leu2Δ1 his3Δ200 trp1Δ63 VPH1-3XmCherry-kanMX6 atg18Δ::hphNT2 ATG18-mNeongreen-hphNT2 leu2Δ1::LEU2-fab1-ha* | This study |
| MUK110 | MATa *ura3-52 leu2Δ1 his3Δ200 trp1Δ63 VPH1-3XmCherry-kanMX6 ATG18-mNeongreen-hphNT2 vac14Δ::6HAklTRP leu2Δ1::LEU2-fab1-ha* | This study |
| MUK111 | MATa *ura3-52 leu2Δ1 his3Δ200 trp1Δ63 VPH1-3XmCherry-kanMX6 atg18Δ::hphNT2 ATG18-FGGG-mNeongreen-hphNT2 leu2Δ1::LEU2-fab1-* | This study |
| MUK113 | MATa *ura3-52 leu2Δ1 his3Δ200 trp1Δ63 VPH1-3XmCherry-kanMX6 ATG18-mNeongreen-hphNT2 vac7Δ::6HAklTRP leu2Δ1::LEU2-fab1-ha* | This study |
| MUK116 | MATa *ura3-52 leu2Δ1 his3Δ200 trp1Δ63 ura3-52::URA3-mCherry-TUB1 ATG18-mNeongreen-hphNT2* | This study |
| MUK117 | MATa *ura3-52 leu2Δ1 his3Δ200 trp1Δ63 VAC7-GFP-His3MX6 VPH1-3XmCherry-natNT2* | This study |
| SEY033 | MATa *ura3-52 leu2Δ1 his3Δ200 trp1Δ63 VPH1-GFP-His3MX6* | This study |
| SEY081 | MATa *ura3-52 leu2Δ1 his3Δ200 trp1Δ63 VPH1-GFP-His3MX6 atg18Δ::hphNT2* | This study |
| SEY082 | MATa *ura3-52 leu2Δ1 his3Δ200 trp1Δ63 VPH1-3XmCherry-His3MX6 ATG18-GFP-klTRP* | This study |
| SEY084 | MATa *ura3-52 leu2Δ1 his3Δ200 trp1Δ63 vac7Δ::His3MX6* | This study |
| SEY213 | MATa *ura3-52 leu2Δ1 his3Δ200 trp1Δ63 vac7Δ::His3MX6 leu2Δ1::LEU2-fab1-ha* | Huda *et al*, 2023 |
| SEY250 | MATa *ura3-52 leu2Δ1 his3Δ200 trp1Δ63 leu2Δ1::LEU2-fab1-ha* | Huda *et al*, 2023 |
| SEY256 | MATa *ura3-52 leu2Δ1 his3Δ200 trp1Δ63 ura3-52::URA3-GFP-TUB1 VPH1-3XmCherry-kanMX6 vac7Δ::6HAklTRP* | Huda *et al*, 2023 |
| SGY008 | MATa *ura3-52 leu2Δ1 his3Δ200 trp1Δ63 VPH1-3XmCherry-kanMX6 FAB1-mNeongreen-hphNT2* | This study |
| SGY009 | MATa *ura3-52 leu2Δ1 his3Δ200 trp1Δ63 VPH1-3XmCherry-kanMX6 VAC14-mNeongreen-hphNT2* | This study |
| MHY334 | MATa ura3-52 *leu2Δ1: his3Δ200 trp1Δ63 VPH1-3xmCherry-kanMX6 SnxA-GFP-URA pho85Δ::klTRP1* | This study |

| | | |
|---|---|---|
| MHY333 | MATa *ura3-52 leu2∆1::LEU2 his3∆200 VPH1-3xmCherry-kanMX6 Snxa-GFP-URA* | This study |

| Plasmid name | Description | Reference |
|---|---|---|
| CGB8.15 | *pHO-KanMX-TEFprom-CPYlead-v-SEP-mCherry-CYC1term* | Okreglak *et al,* 2023 |
| pAFS125-GFP-TUB1 | *GFP-TUB1* in a *URA3* -based integration vector | Straight *et al,* 1997 |
| pAK011 | *mCherry-TUB1* in a *URA3* -based integration vector | Khmelinskii et al, 2000 |
| pCD001 | pRS315 *Atg18-EGFP-klTRP* | This study |
| pCD002 | pRS315 *Atg18Sloop-EGFP-klTRP* | This study |
| pCD003 | pRS315 *Atg18FGGG-EGFP-klTRP* | This study |
| pMH007 | pRS306-*TEF-SNXA-GFP-Cyc-term.* | This study |
| pMH009 | pRS306-*TEF-SNXA-mCherry-Cyc-term.* | This study |
| pMH011 | pRS304-*TEF-SNXA-mCherry-Cyc-term.* | This study |
| pOK104 | pRS416*TEF-EGFP-2xFYVE-URA* | Gillooly *et al* , 2000; Obara *et al* , 2008 |
| pSE003 | pRS305-*fab1* ( T2250A, E1822V, F1833L) | Huda *et al,* 2023 |
